# A Comparative Benchmark of Biomedical Language Models for Concept Normalization from Real-World Text

**DOI:** 10.64898/2026.09.19.752843

**Authors:** Anshul Verma, Abhijay, Manan Vangani, Satyartha Prakash, Kumardeep Chaudhary

**Affiliations:** CSIR-Institute of Genomics and Integrative Biology, New Delhi, 110007 India; Academy of Scientific and Innovative Research (AcSIR), Ghaziabad, 201002 India; Amity Institute of Biotechnology, Amity University, Noida, 201313 India

**Keywords:** Medical Concept Normalization, BERT, LLMs, UMLS, SNOMED CT, MedDRA, SapBERT, Qwen 2 Instruct

## Abstract

Patient-reported and clinically documented narratives often contain informal, fragmented and linguistically heterogeneous expressions, complicating medical concept normalization (MCN) and frequently necessitating pre-processing before terminology mapping. Despite rapid advances in biomedical language modeling, the comparative utility of representation models for MCN and their integration with instruction-tuned LLMs in automated normalization workflows remain underexplored. In this study, we systematically benchmarked 15 general, biomedical and clinical transformer-based representation models together with 12 instruction-tuned LLMs across 12,713 instances from five established datasets: TAC2017_ADR, TwADR-L, TwiMed, CADEC and SMM4H2017. Representation models were evaluated using embedding-based semantic retrieval, whereas instruction-tuned LLMs were assessed as upstream text-correction modules. SapBERT achieved the highest Top-5 accuracy among representation models, reaching 63.8% for SNOMED CT and 58.0% for MedDRA without correction. Qwen 2 Instruct was selected as the preferred corrector on the basis of its favorable balance between Top-1 normalization performance and computational efficiency relative to substantially larger models, including Llama 3.1 Instruct (70B). Incorporation of Qwen 2 Instruct increased Top-5 accuracy to 69.4% for SNOMED CT and 63.2% for MedDRA. The resulting framework accepts heterogeneous short medical expressions without manual input pre-processing and automatically performs text refinement, semantic retrieval and terminology mapping to standardized concepts and vocabulary codes. These findings establish a benchmark-guided, scalable framework for automated medical terminology standardization.

## Introduction

The dialogue between a patient and a healthcare practitioner is deeply insightful regarding several aspects such as the patient’s understanding of their own health including signs and symptoms of a disease/disorder. This interaction, however, is found to be a constant to and fro of lay phrases from the patient and medical lingo from the clinician. A similar pattern appears on social media platforms as well in the form of posts or under threads on medical forums where a global populace reports their experience ranging from adverse drug reactions to side effects of a drug or personal experiences through a disease/disorder. Medical Concept Normalization (MCN) therefore aims to map such heterogeneous medical language to globally standardized biomedical concepts maintained in large ontologies such as the Unified Medical Language System (UMLS), SNOMED CT and MeSH, supporting terminology harmonization and downstream medical applications [1–5]. This task is particularly challenging for patient-generated and social media texts which are mostly informally phrased, ambiguous and vague in symptomatic descriptions [6, 7].

From an historical perspective, MCN has seen significant advancement over the years, from rudimentary deterministic string matching and rule-based approaches to machine learning and deep learning models capable of handling noisy and unstructured text [8–10]. Encoder-based models or transformers subsequently introduced contextual awareness in text which was a huge leap from feature dependence [11–13]. Two-stage pipelines that implemented sparse candidate generators (e.g., BM25) along with neural re-rankers further improved retrieval [14, 15], while metric learning approaches such as SapBERT and BioSyn have proven their through with optimized biomedical synonym alignment in embedding space [16–18]. Graph-based pipelines and encoder-decoder architectures have also been explored of late to infuse ontological and/or structured knowledge and contextual reasoning into alignment tasks [19–22]. Despite this, most pipelines and workflows are constrained by upstream components such as Named Entity Recognition (NER), valid and curated annotations and highly domain-specific vocabularies [3, 23]. These shortcomings limit their real-world applications in normalizing noisy and unstructured text. Duplication, class imbalance and unintentional incomplete data coverage can additionally inflate apparent performance and limit generalizability [18, 23–25]. As a result, there is a growing need for ontology-scale, concept-centric normalization frameworks that minimize pre-processing requirements.

These limitations are amplified in noisy, free-flowing and unstructured text. Rudimentary approaches rely on pre-processing, explicit mention extraction and n-gram-based methods to improve contextual representation beyond classical NER and annotation [26]. However, serializing NER and MCN remains difficult because errors in entity detection propagate directly to normalization [27]. Several frameworks integrate entity recognition and normalization, but each presents important limitations. MetaMap provides UMLS mapping but faces challenges with spelling variations and unclear concept mentions [28]; Bio-YODIE improves disambiguation but requires annotated data [29]; cTAKES uses a modular NLP pipeline that may require additional domain-specific components [30]; ScispaCy combines supervised NER with a string-matching architecture [31] and MedCAT generally benefits from annotated data for optimal performance [32]. More recent open-source approaches such as HunFlair2, preon, Thera-Py and xMEN extend neural, multilayer, graph-based or multilingual generate-and-rank normalization, but introduce dependencies on pretrained resources, domain-specific design or additional architectural complexity [33–36].

Large language models (LLMs) offer substantial potential for clinical free-text processing, but their stochasticity limits reliable structured retrieval. Klang et al. concluded that LLMs are unsuitable as standalone tools for structured clinical data tasks without explicit computational scaffolding [37]. This limitation is particularly important for ontology-scale concept normalization across UMLS, where the search space comprises millions of concepts and precise retrieval is essential. In line with this, this study positions instruction-tuned LLMs strictly as upstream correctors for informal text while retaining the vectorization space as the driver of normalization and retrieval.

Accordingly, we develop an end-to-end framework that incurs no pre-processing steps such as manual annotation, entity extraction and/or adherence to strict grammar norms on the user’s end. The seamless pipeline takes patient-generated free-form text as input to be subsequently normalized and mapped to standardized medical vocabularies and the output is populated with several features such as normalized medical terms, parent vocabulary codes and cosine similarity scores, with no human intervention required apart from the query itself. Another strength of the study lies in the vast latent space serving as the foundational store of vectors, which is not limited to any one lexicon dataset or a specific medical vocabulary but represents the whole repository of vocabularies recorded under the UMLS [38]. Each concept is enriched with its respective aliases/synonyms recorded against it in the UMLS. The latent space is deeper and encodes more knowledge as we opt for a term-level mapping rather than concept-level mapping alone.

## Methods and Materials

### Dataset

For constructing the semantic embedding spaces across multiple transformer-based models, we employed the Unified Medical Language System (UMLS; version 2025AA) as a foundational, standardized repository of biomedical vocabularies and corresponding concept identifiers. UMLS delivers an integrative metathesaurus that harmonizes a wide spectrum of authoritative, clinical and biomedical vocabularies including but not limited to the Human Phenotype Ontology (HPO), International Classification of Diseases (ICD), Medical Dictionary for Regulatory Activities (MedDRA) and Systematized Nomenclature of Medicine-Clinical Terms (SNOMED CT) **(Supplementary Table S1)**. The UMLS 2025AA release statistics record ∼3.4 million concepts compiled from 190 different sources. This comprehensive integration enables unified semantic representation and cross-terminology mapping, thereby providing a robust substrate for large-scale MCN and retrieval tasks.

### Data Pre-processing

Following the approval of our request for the access to the UMLS Metathesaurus License, the task of data pre-processing was performed locally leveraging the MetamorphoSys tool. After sub-setting the vast archive of UMLS for only the pertinent English standard medical vocabularies, we obtained a set of 40 Rich Release Format (RRF) files. All the files serve as a connected database with the UMLS Concept Unique Identifier (CUI) acting as the common key. The central file MRCONSO (stores names, synonyms), enriched with information from MRDEF (stores definitions), MRSAB (stores source abbreviations) and MRSTY (stores semantic types), was selected to serve as a comprehensive knowledge base.

Coarsely representing every point of data with all its features in the embedding space would be exhaustive in terms of data coverage but the resulting latent space would be dense and noisy, ultimately compromising retrieval performance. As a trade-off, only the preferred names and the related synonyms and terms concerning those concepts are mapped onto the latent space. Features such as the UMLS CUIs, source vocabulary identifiers and definitions are utilized to populate a rich metadata store.

### Transformer-based Models for Embedding Space

To develop a robust semantic embedding space for concept normalization tasks, fifteen distinct transformer-based language models were systematically benchmarked. These models were selected to represent a diverse spectrum of pre-training corpora and architectural objectives, encompassing general-domain, biomedical and clinical language models (Table 1). Each model captures distinctive linguistic and semantic characteristics due to its heterogeneous training corpora and architectural objectives. This diversity calls for a comparative assessment of their capacity to encode biomedical semantics and their capabilities to support accurate normalization through embedding-based similarity mapping.

**Table 1:** Summary of the fifteen transformer-based language models evaluated for MCN with their pre-training corpus.

| Model | Year | Training Corpus | Parameters |
| --- | --- | --- | --- |
| BioClinicalBERT [39] | 2019 | MIMIC-III clinical notes | 110M |
| BioBERT [40] | 2019 | English Wikipedia + BooksCorpus + PubMed abstracts + PMC full-text articles | 110M |
| ClinicalBERT [13] | 2019 | MIMIC-III clinical notes | 110M |
| GPT2-large [41] | 2019 | WebText | 774M |
| BioM-ALBERT [42] | 2021 | PubMed + PMC full-text articles | 223M |
| BioM-ELECTRA [42] | 2021 | PubMed abstracts | 110M |
| BioMedBERT [12] | 2021 | PubMed abstracts + PMC full-text articles | 110M |
| SapBERT [16] | 2021 | PubMed + PMC full-text articles + UMLS pairs + self-alignment training | 110M |
| UmlsBERT [43] | 2021 | MIMIC-III clinical notes + UMLS concepts | 110M |
| BioLinkBERT [44] | 2022 | PubMed abstracts with explicit citation link information | 110M |
| CODER [45] | 2022 | UMLS terms + Relation triplets | 110M |
| MedBERT [46] | 2022 | BioNLP + CRAFT + N2C2 + Biomedical Wikipedia texts | 110M |
| BioGPT [47] | 2023 | PubMed abstracts | 350M |
| ModernBERT [48] | 2024 | Web + code + scientific literature | 120M |
| BioClinical ModernBERT [49] | 2025 | PubMed abstracts + PMC full-text articles + MIMIC-III clinical notes + MIMIC-IV clinical notes + CheXpert Plus + INSPECT + ADE Corpus + Social History notes, Chief complaints, Pathology reports, Stroke MRI reports, Simulated interviews and others | 120M |

A few models with the primary objective of architectural optimization of the original BERT family, namely BioM-ALBERT, BioM-ELECTRA, ModernBERT and BioClinical ModernBERT are also evaluated against earlier architectures to reflect current research efforts and improvements in the study. Two open-source autoregressive models, GPT2-large and BioGPT, were also evaluated to ensure a comprehensive battery of models.

### Instruct Large Language Models as Biomedical Correctors

The key issue in MCN is the innate unstructured nature of patient-reported symptoms. These colloquial expressions often consist of non-standard medical terms, misspellings and vague descriptions that result in semantic mismatching when processed by transformer-based encoders. To address this issue, we evaluated twelve LLMs as upstream biomedical correctors (Table 2).

**Table 2:**
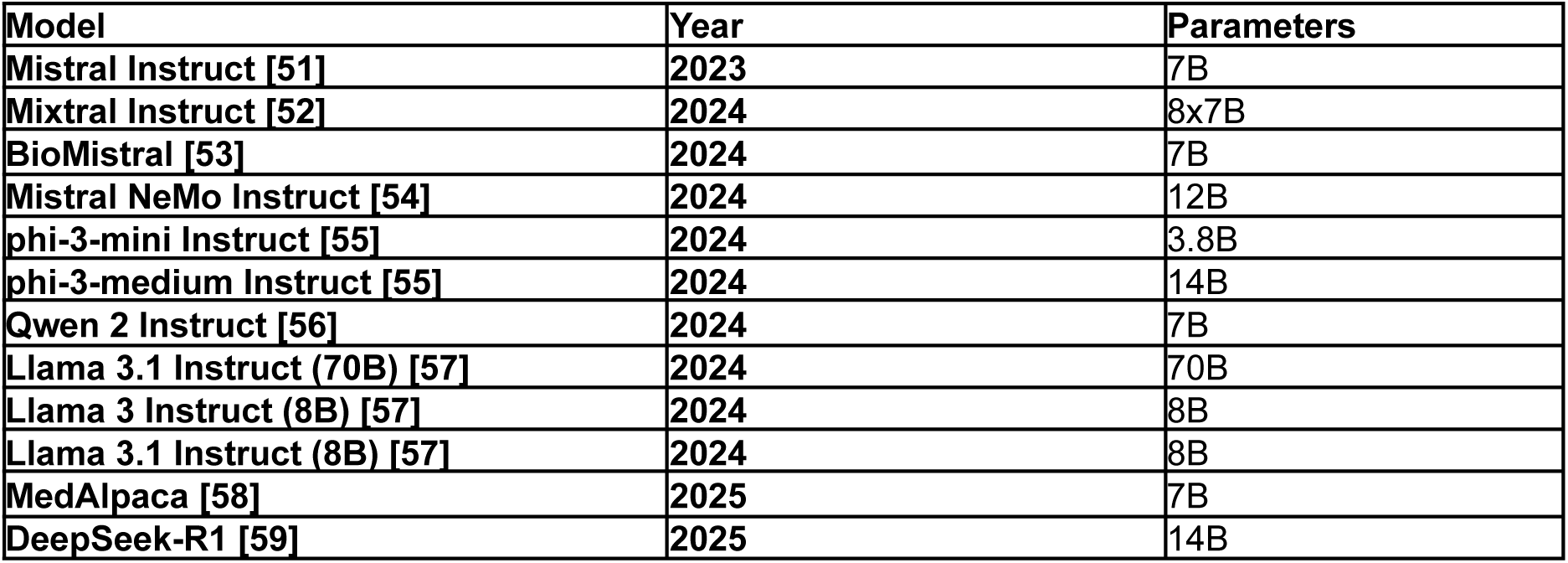
Summary of the twelve LLMs evaluated as biomedical correctors.

We developed four distinct prompts (Supplementary Figure S1), each designed to test different instruction strategies and used to instruct the LLMs to rephrase raw, loose-text entries into relevant biomedical text avoiding any diagnostic speculation [50]. The objective was to refine the input signal to better map with the term-level embedding space and fix spelling errors that result in erroneous normalization while mitigating the risk of “semantic drift”, where a patient’s actual symptom text is incorrectly mapped to a similar but clinically unrelated disorder term.

### Pipeline Architecture

To develop the end-to-end MCN pipeline, we designed a multi-stage architecture that systematically transforms raw, unstructured free-text inputs into standardized medical concepts with selected ontology identifiers (Figure 1).

**Figure 1:**
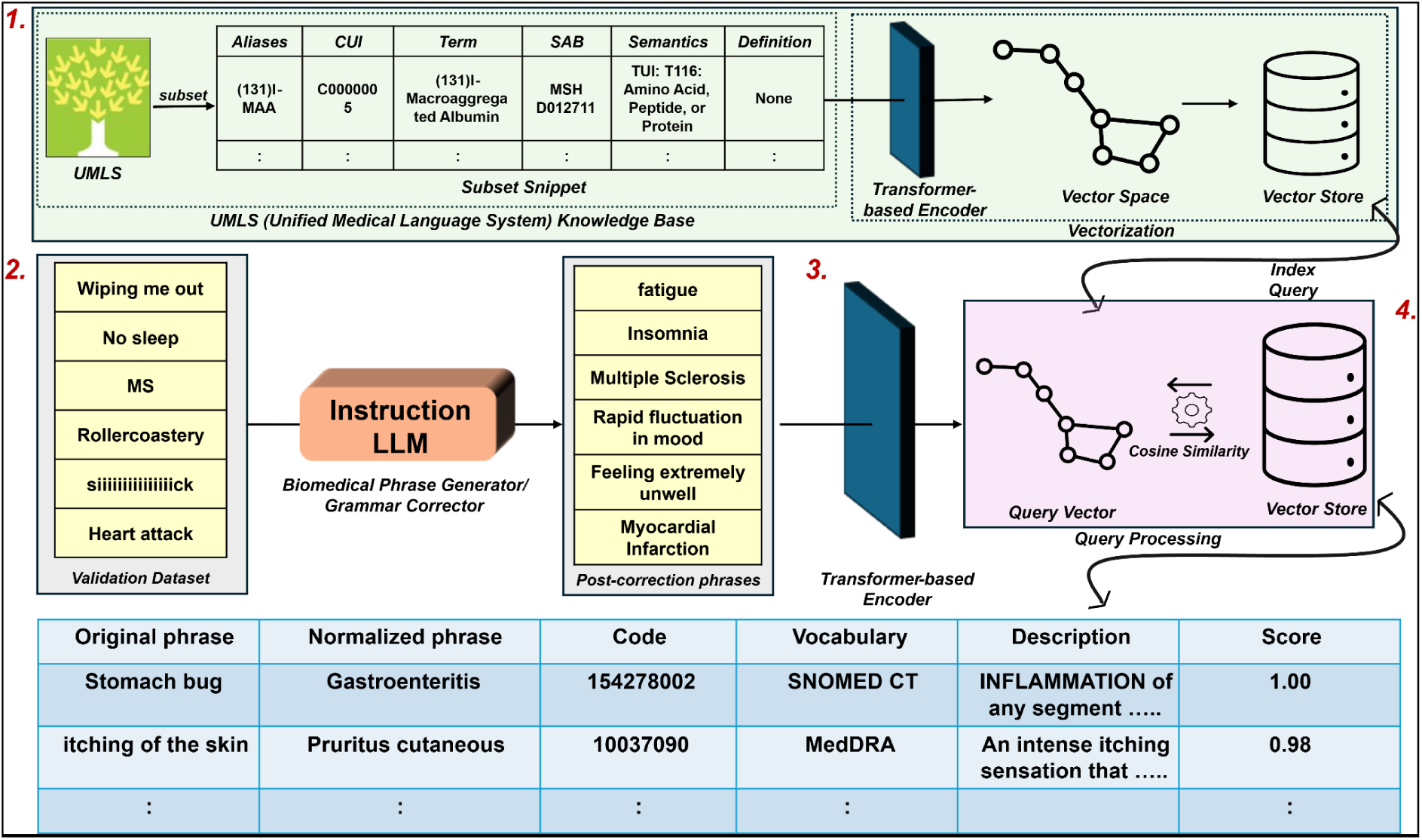
Schematic of the MCN pipeline, illustrating free-text correction, embedding-based semantic matching against the UMLS concept space and retrieval of standardized medical terms with corresponding ontology codes.

In the first stage, the raw query input instance is fed to an instruct LLM-based biomedical corrector which uses a specialized prompt to convert a loose-text phrase into a relevant biomedical phrase. By doing so, many of the instances that were earlier incorrectly mapped in the absence of an upstream corrector were correctly normalized to the relevant medical terminology.

In the second stage, the corrected query instance is then encoded into a dense semantic representation using each of the fifteen transformer-based language models. These models, which differ in pre-training corpora, architectural design and optimization objectives, generate embeddings that capture varying degrees of semantic understanding, enabling a comprehensive assessment of model-specific performance in representing medical queries.

In the subsequent stage, the query embedding is matched against the pre-calculated embedding space consisting of all UMLS terms using cosine similarity score as the similarity metric. Prior to indexing and retrieval, both query and term embeddings were subjected to L2 normalization, enabling cosine similarity computation through inner-product search. For efficient and scalable nearest-neighbor retrieval over the large UMLS embedding space, we employ Facebook AI Similarity Search (FAISS) [60] using the IndexFlatIP indexing strategy, which performs exact brute-force inner-product search without approximation. Embedding dimensionality was inferred dynamically at runtime, enabling compatibility across heterogeneous transformer architectures.FAISS supports high-performance similarity search through optimized indexing structures, allowing rapid identification of the most semantically similar concept embeddings. The nearest-neighbor with the highest cosine similarity score is selected as the most probable normalized representation of the input.

In the final stage, all the retrieved embeddings are mapped back to their corresponding vocabulary identifier along with their standardized medical terms followed by deduplication of redundant identifiers. Finally, the deduplicated candidates are ranked by cosine similarity score and returned as the system output. This ensures that users receive not only a standardized medical phrase but also its precise ontology-grounded code, facilitating interoperability, downstream analytics and clinical decision support.

### MedNorm Dataset (Evaluation Dataset)

The performance of the proposed MCN pipeline was evaluated using MedNorm [61], a publicly available benchmark corpus comprising 27,979 textual phrases annotated with standardized medical concept codes. MedNorm provides mappings to MedDRA and SNOMED CT codes and integrates data from five datasets spanning biomedical literature and social media domains: TAC2017_ADR, TwADR-L, TwiMed, CADEC and SMM4H2017. These sources collectively encompass a wide range of linguistic variability including informal patient-generated content, clinically relevant adverse event descriptions and domain-specific biomedical expressions.

These datasets were treated as blind evaluation datasets and no instances from this corpus were used during embedding construction or model selection, ensuring an unbiased assessment of generalization performance. Each record contains a raw textual phrase along with curated annotations including the corresponding UMLS CUI and best-mapped SNOMED CT and MedDRA codes. Model-mapped annotations were compared against these reference annotations using exact-match criteria to assess normalization performance.

### Quality Issues with the Blind Dataset

The MedNorm corpus aggregates multiple source datasets that were originally annotated with different terminology systems and not all datasets contained complete mapping to UMLS, SNOMED CT and MedDRA at the time of their original release. Although the MedNorm resource subsequently provides harmonized mappings across these terminologies, the availability and completeness of identifiers vary across constituent datasets [61]. The distribution of terminology mappings across datasets, including the number of instances linked to UMLS, SNOMED CT and MedDRA, is summarized in Supplementary Table S2. For the present study, SNOMED CT- and MedDRA-linked instances were selected to enable consistent concept-level evaluation.

During pre-processing, we identified redundant textual instances within and across dataset segments. Exact duplicates were removed across datasets, followed by an additional case-insensitive deduplication step to eliminate entries differing only by letter case. After removing all redundant instances within each dataset, the final blind evaluation dataset consisted of 12,713 unique textual descriptions, which were used for all downstream normalization experiments (Table 3).

**Table 3:** Overview of the MedNorm evaluation dataset used for blind evaluation, summarizing phrase counts and corresponding SNOMED CT, MedDRA and UMLS CUI identifiers across constituent datasets after eliminating redundancy.

| Dataset | Phrases | SNOMED IDs | MedDRA IDs | UMLS CUIs |
| --- | --- | --- | --- | --- |
| TAC2017 ADR | 2,106 | 2,106 | 2,106 | 0 |
| TwADR-L | 2,581 | 2,581 | 2,581 | 2,581 |
| Twimed | 864 | 864 | 864 | 864 |
| CADEC | 3,376 | 3,376 | 3,376 | 0 |
| SMM4H2017 | 3,786 | 3,786 | 3,786 | 0 |
| Total | 12,713 |  |  | 3,445 |

### Statistics and Programming Environment Statistical Performance Evaluation

Model performances were quantitatively assessed using standard information retrieval and classification metrics appropriate for MCN tasks. Accuracy, as the primary evaluation metric, was computed by comparing the retrieved normalized concept against the ground-truth annotations provided in the blind evaluation dataset (segmented MedNorm). A mapping was considered correct when the retrieved concept identifier exactly matched the reference identifier (likewise for Top-K).

To further assess retrieval quality, we also computed Mean Reciprocal Rank (MRR), Recall and Normalized Discounted Cumulative Gain (nDCG). These metrics evaluate the ranking of the correct concept among retrieved candidates and provide a more comprehensive assessment of normalization performance. Accordingly, overall accuracy, accuracy@K, Recall@K, MRR@K and nDCG@K were reported for all evaluated models to enable consistent comparison across the benchmarking framework. Reproducibility of both transformer-based models and instruct LLMs was additionally assessed across repeated randomized input-order runs using observation-level stability, pairwise agreement, Cohen’s κ, Cochran’s Q, pairwise McNemar testing and practical-equivalence analysis.

### High-Performance Computing and GPU Configuration

All computational workflows involved in embedding generation, query normalization and concept retrieval were executed on high-performance computing (HPC) infrastructure with GPU acceleration. The pipeline was implemented in Python v3.10.18 using PyTorch v2.3.0 with CUDA 12.2 enabled.

A single NVIDIA A100-SXM4 GPU with 80GB memory was utilized for large-scale embedding computation across all evaluated instruct-based and transformer-based language models.

## Results and Discussion

The proposed MCN pipeline was evaluated on the blind datasets comprising 12,713 textual instances in total, annotated with standardized medical concepts. Evaluation was conducted in two phases. In the initial phase, overall and Top-K accuracies were calculated across all datasets for SNOMED CT and MedDRA as gold-standard reference codes for each model. In the subsequent phase, the best-performing transformer model was supplemented with an LLM-based correction module upstream. All LLM-based correction models were prompted using a specialized set of instructions shown in Figure 2.

**Figure 2:**
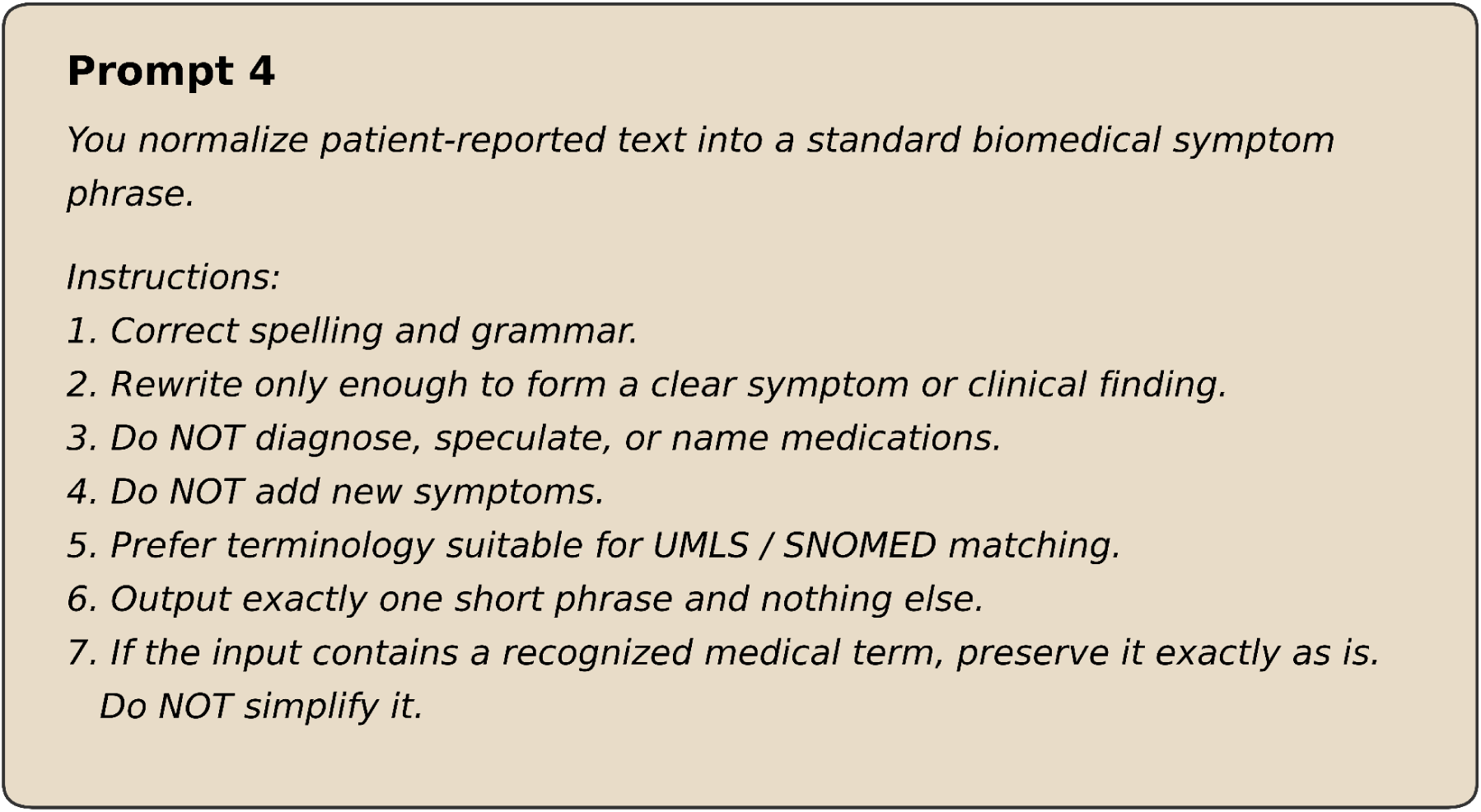
Prompt template for query correction.

As described in Methods and Materials, four distinct prompts were initially designed manually to assess different sets of instructions for correcting identical instances from the blind evaluation dataset (SNOMED CT). For final benchmarking, considering the time constraints associated with testing four prompts against 12,713 queries for each of the 12 instruct models, prompt-selection was optimally performed on a subset of the validation dataset containing particularly challenging instances (n = 1,493) that were incorrectly mapped by all 15 of the transformer-based models in phase 1. The best-performing prompt (Prompt 4) as per overall accuracy was thereafter chosen for all LLM-based correction experiments. The intersection of instances correctly normalized by no model (n = 1,493) is shown in Supplementary Figure S2, while comparative prompt benchmarking results are shown in Supplementary Figure S3.

### Benchmark Evaluation of Transformer-based Models

Figure 3 shows dataset-wise full-retrieval accuracy across all the transformer-based models further depicting substantial heterogeneity in normalization performance when SNOMED CT codes are the ground-truth. In Figure 3A, SapBERT clearly outperforms all other models with a full-retrieval accuracy of 96.3% for TAC2017_ADR, 85.9% for TwADR-L, 86.2% for TwiMed, 86.6% for CADEC and 68.3% for SMM4H2017. SapBERT is closely followed by CODER, demonstrating effectiveness of contrastive learning to pull synonymous medical concepts together within a similar semantic space.

**Figure 3:**
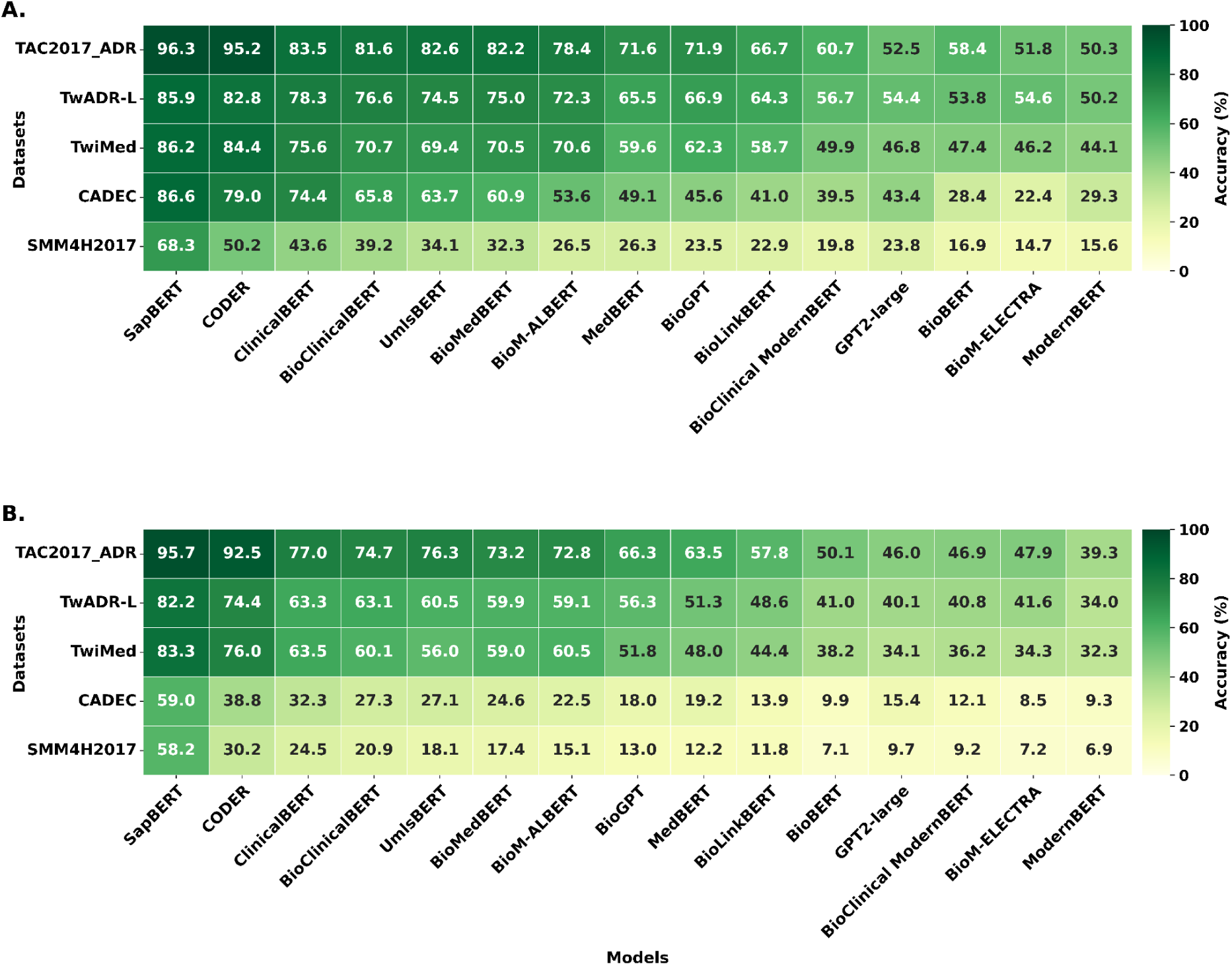
Normalization performance across transformer-based models evaluated on segmented MedNorm corpus with models ordered by mean full-retrieval accuracy across dataset segments. **A)** SNOMED CT-based evaluation **B)** MedDRA-based evaluation.

Domain-specific models such as ClinicalBERT, BioClinicalBERT, UmlsBERT, BioMedBERT, MedBERT, BioLinkBERT and BioBERT lack such a contrastive training objective and therefore span a wide range of accuracies (16.9 - 83.5%). Although UmlsBERT incorporates an additional embedding dimension of UMLS semantic types of concepts as an architectural enhancement, the results still show that it still struggles to align synonymous concepts.

BioM-ALBERT and BioM-ELECTRA (architecturally optimized for masked language modeling), along with the autoregressive models BioGPT and GPT2-large, achieve below-average to low accuracies (14.7 - 78.4%). These standings may also be explained by a difference in the training objectives of the autoregressive models which are tasked with predicting the next word rather than optimizing vector similarity.

The most recent models, ModernBERT and BioClinical ModernBERT, trained on web-scale data as well as large and diverse biomedical corpora, were also clearly outperformed by SapBERT, with results ranging from 15.6% to 60.7%. Despite being trained on broader data and using updated architectural improvements, these models remain relatively general-purpose and were not explicitly optimized to cluster semantically synonymous biomedical terms in the embedding space.

It is also notable that SMM4H2017 is the most challenging dataset regardless of the transformer-based embedding model. The most imminent reason being that this dataset majorly consists of idiomatic, vague and slang-heavy colloquial phrases such as “*feel like death*”, “*running around like a mad man*”, “*into an emotionless zombie*” etc.

To extend the evaluation, Supplementary Figure S4A reports Top-K accuracies (K=1, 5, 10, 15, 20, 30, 40, 50) for the same five datasets across all the models. The performance was broadly preserved with the same trend where SapBERT and CODER consistently outperform all other models for all datasets. Notably, a plateau in performance for TAC2017_ADR, TwADR-L and TwiMed is seen at Top-K=20,, suggesting a practical threshold after which marginal gains are achieved for the retrieval of correct medical codes by most of the models whereas for CADEC and SMM4H2017 retrieval still improves with depth. SapBERT gains and maintains an impressive lead for SMM4H2017 starting from K=5.

To provide a comprehensive assessment of retrieval quality, additional ranking-based metrics including Recall@K, MRR@K and nDCG@K were also evaluated (Supplementary Figure S5). These analyses demonstrated trends consistent with the primary accuracy results, with SapBERT and CODER consistently achieving the highest retrieval effectiveness across datasets, thereby confirming that the observed performance hierarchy was not dependent on Top-1 accuracy as the sole performance indicator.

For SNOMED CT, statistical comparison confirmed significant differences across the 15 transformer models at all retrieval depths (Cochran’s Q, P<0.001). SapBERT achieved the highest full-retrieval accuracy of 82.6% (95% CI 81.94 – 83.25%), followed by CODER at 74.25%. After Holm correction, 103 of 105 pairwise comparisons remained significant (Supplementary Tables S3-S6).

For MedDRA-based normalization (Figure 3B), SapBERT consistently achieved the highest full-retrieval accuracy again across all five benchmark datasets, reaching 95.7% on TAC2017_ADR, 82.2% on TwADR-L, 83.3% on TwiMed, 59.0% on CADEC and 58.2% on SMM4H2017. CODER ranked second, with corresponding accuracies of 92.5%, 74.4%, 76.0%, 38.8% and 30.2%, respectively. Other domain-specific models, including ClinicalBERT, BioClinicalBERT, UmlsBERT and BioMedBERT, showed moderate performance but remained substantially below SapBERT, particularly on CADEC and SMM4H2017. BioM-ELECTRA and ModernBERT were among the weakest performers, with accuracies declining to 8.5% and 9.3% on CADEC and 7.2% and 6.9% on SMM4H2017, respectively.

In the Top-K retrieval analysis (Supplementary Figure S4B), the same overall model hierarchy was retained across datasets, with SapBERT and CODER consistently showing the strongest performance as K increased. The largest gains were generally observed between K=1 and K=10, followed by a gradual plateau, while CADEC and SMM4H2017 remained the most challenging datasets.

The ranking-sensitive metrics (Supplementary Figure S6) further supported the trend. Across Recall@K, MRR@K and nDCG@K, SapBERT consistently achieved the highest scores, followed by CODER, confirming that its superior performance was maintained beyond Top-1 accuracy and across deeper retrieval thresholds. For MedDRA, model performance also differed significantly across retrieval depths (Cochran’s Q, P<0.001). SapBERT again ranked first with 71.19% full-retrieval accuracy, followed by CODER at 54.89% with 103 of 105 pairwise comparisons significant after Holm correction (Supplementary Tables S7-S10).

### Percentage-Point Gains Achieved from Top-1 to Top-50 Retrieval

Figure 4 compares Top-1 exact-match accuracy with the additional percentage-point gain achieved when retrieval is expanded to the Top-50 candidates. Figure 4A summarizes the average and dataset-specific gains for SNOMED CT normalization. BioGPT showed the largest average improvement (+31 percentage points), followed closely by SapBERT (+30), GPT2-large (+27), MedBERT (+26) and CODER (+25). Dataset-wise, the largest gains were observed for BioGPT on TAC2017_ADR (+41), TwADR-L (+34) and TwiMed (+33), whereas SapBERT showed the greatest improvement on CADEC (+37) and SMM4H2017 (+41).

**Figure 4:**
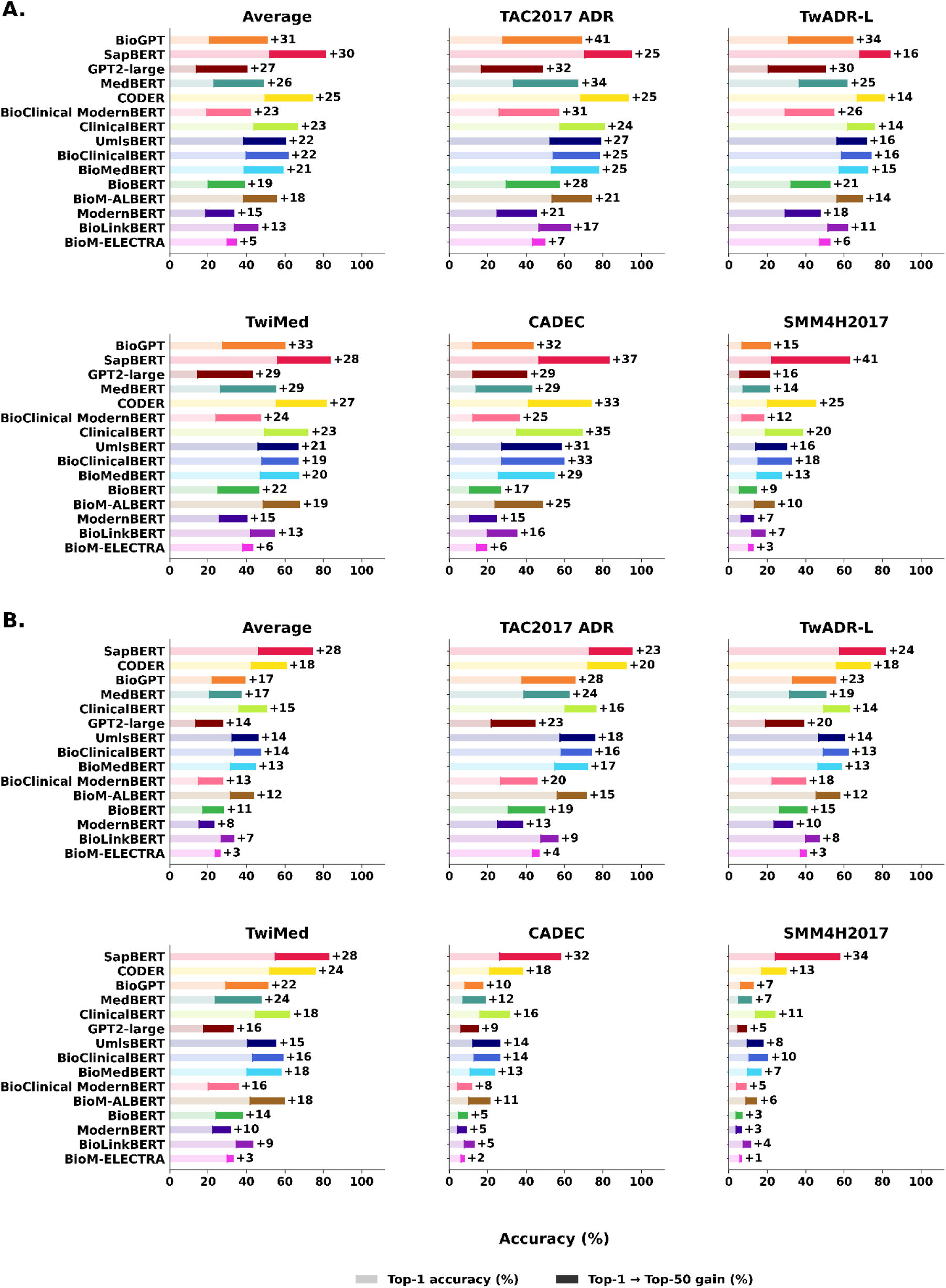
Gains in retrieval accuracy obtained by expanding the candidate set from Top-1 to Top-50. Each panel shows the Top-1 accuracy and the additional percentage-point gain achieved at Top-50 across benchmark datasets for **A)** SNOMED CT and **B)** MedDRA normalization. Larger gain segments indicate greater recovery of the correct concept at increased retrieval depth.

Figure 4B presents the corresponding analysis for MedDRA normalization. SapBERT demonstrated the largest average Top-1-to-Top-50 gain (+28 percentage points), followed by CODER (+18), BioGPT and MedBERT (+17 each). SapBERT also showed the greatest improvement on TwADR-L (+24), TwiMed (+28), CADEC (+32) and SMM4H2017 (+34), while BioGPT exhibited the largest gains on TAC2017_ADR (+28). Overall, the results indicate that increasing retrieval depth substantially improves concept recovery.

### Comparative Analysis of Instruction-Tuned LLMs as Biomedical Correctors

The incorporation of instruction-tuned LLMs as a correction layer significantly enhanced the SapBERT-based normalization pipeline’s performance, which serves here as the baseline. For SNOMED CT normalization (Figure 5A), Llama 3.1 Instruct (70B) achieved the strongest overall performance, with consistently high full-retrieval accuracy across all five datasets, including TAC2017_ADR (95.0%), TwADR-L (88.0%), TwiMed (89.9%), CADEC (89.7%) and SMM4H2017 (82.2%). Qwen 2 Instruct showed closely comparable performance, particularly on TAC2017_ADR (95.0%), TwADR-L (87.8%), TwiMed (89.3%) and CADEC (89.1%). The largest improvements were observed for linguistically difficult datasets such as SMM4H2017, likely because instruction-tuned models can transform slang-heavy, misspelled and fragmented expressions into terminology that is semantically closer to standardized medical concepts. In contrast, gains were smaller for relatively structured datasets such as TAC2017_ADR, where unnecessary reformulation may introduce semantic drift and move already precise expressions away from the reference terminology. The phi-3-medium Instruct remained the weakest corrector across datasets, indicating that correction quality depends not only on model size but also on the ability to preserve clinical meaning during reformulation.

**Figure 5:**
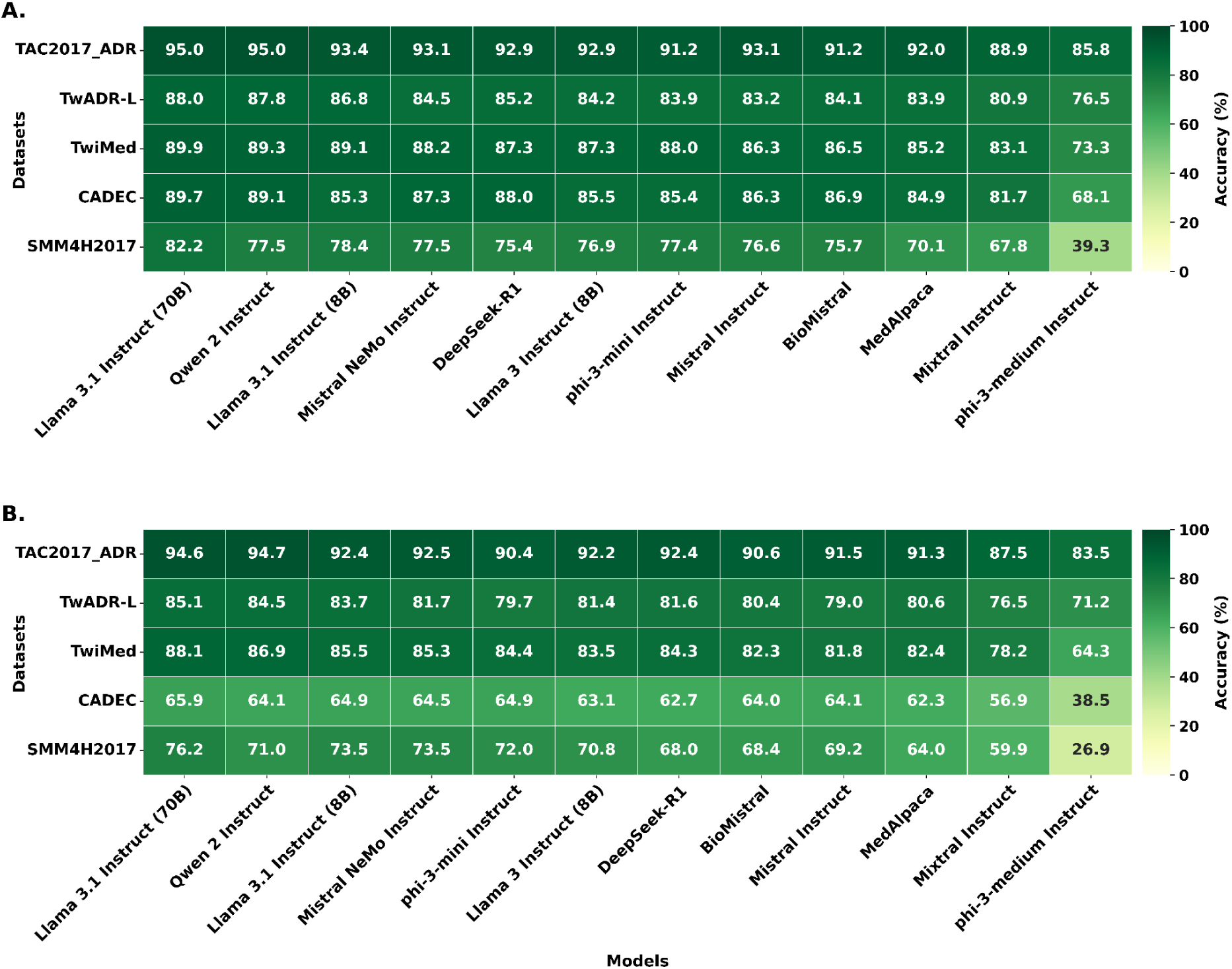
Normalization performance of SapBERT when appended with Instruct models (as corrector), evaluated on segmented MedNorm corpus with models ordered by mean full-retrieval accuracy across dataset segments. **A)** SNOMED CT-based evaluation and **B)** MedDRA-based evaluation.

Supplementary Figure S7A, shows that Top-K retrieval follows the same pattern, with most gains occurring between K=1 and K=10 before reaching a gradual plateau. Llama 3.1 Instruct (70B), Qwen 2 Instruct and Mistral NeMo Instruct consistently occupied the upper performance range, whereas phi-3-medium Instruct remained substantially lower. The complementary ranking metrics in Supplementary Figure S8, further confirm this trend across Recall@K, MRR@K and nDCG@K. Qwen 2 Instruct remained particularly competitive despite its substantially smaller parameter count relative to Llama 3.1 Instruct (70B), suggesting a more favorable balance between retrieval performance and computational demand.

For SNOMED CT, statistical comparison across the 12 instruction-tuned LLMs confirmed significant differences at all retrieval depths (Cochran’s Q, P<0.001). At full-retrieval, Llama 3.1 Instruct (70B) achieved the highest overall accuracy of 88.04% (95% CI 87.49 – 88.61%), followed by Qwen 2 Instruct at 86.36%. After Holm correction, 53 of 66 pairwise comparisons remained significant (Supplementary Tables S11-S14).

For MedDRA normalization (Figure 5B), overall performance was lower than in the SNOMED CT setting, but a similar model hierarchy was observed. Llama 3.1 Instruct (70B) again achieved the strongest and most consistent performance, reaching 94.6% on TAC2017_ADR, 85.1% on TwADR-L, 88.1% on TwiMed, 65.9% on CADEC and 76.2% on SMM4H2017. Qwen 2 Instruct remained highly competitive with accuracies of 94.7%, 84.5%, 86.9%, 64.1% and 71.0% across the respective datasets. Llama 3.1 Instruct (8B) and Mistral NeMo Instruct also maintained comparatively strong performance, whereas phi-3-medium Instruct again showed the greatest degradation, particularly for CADEC and SMM4H2017.

The MedDRA Top-K analysis in Supplementary Figure S7B similarly showed substantial improvement between K=1 and K=10, followed by progressively smaller gains toward K=50. The advantage of stronger corrector models remained evident across retrieval depths, particularly for the more difficult CADEC and SMM4H2017 datasets. Consistently, Supplementary Figure S9 demonstrated that Recall@K, MRR@K and nDCG@K preserved the same broad ranking pattern, with Qwen 2 Instruct and Llama 3.1 Instruct (70B) remaining among the leading models. Together, these results suggest that LLM correction is most beneficial when the input is linguistically noisy, while over-correction of already standardized expressions remains an important source of error.

For MedDRA, performance also differed significantly across the 12 LLMs (Cochran’s Q, P<0.001). Llama 3.1 Instruct (70B) again ranked first with 79.14% full-retrieval accuracy, followed by Llama 3.1 Instruct (8B) at 77.24%, with 58 of 66 pairwise comparisons remaining significant after Holm correction (Supplementary Tables S15-S18).

In Top-1 gain/loss performance for SNOMED CT shown in Figure 6A, most instruction-tuned LLMs improved upon the standalone SapBERT baseline accuracy of 48.3%. Qwen 2 Instruct achieved the largest gain (+6.3 percentage points), followed by Mistral NeMo Instruct (+5.5 pp), Llama 3.1 Instruct (8B) (+4.0 pp) and Llama 3.1 Instruct (70B) (+3.7 pp). In contrast, phi-3-medium Instruct showed the largest degradation (-25.4 pp), followed by Mixtral Instruct (-7.3 pp), phi-3-mini Instruct (-2.2 pp) and Mistral Instruct (-0.8 pp). The poor performance of phi-3-medium may reflect weaker adherence to the constrained correction prompt, including generation of unnecessarily conversational responses.

**Figure 6:**
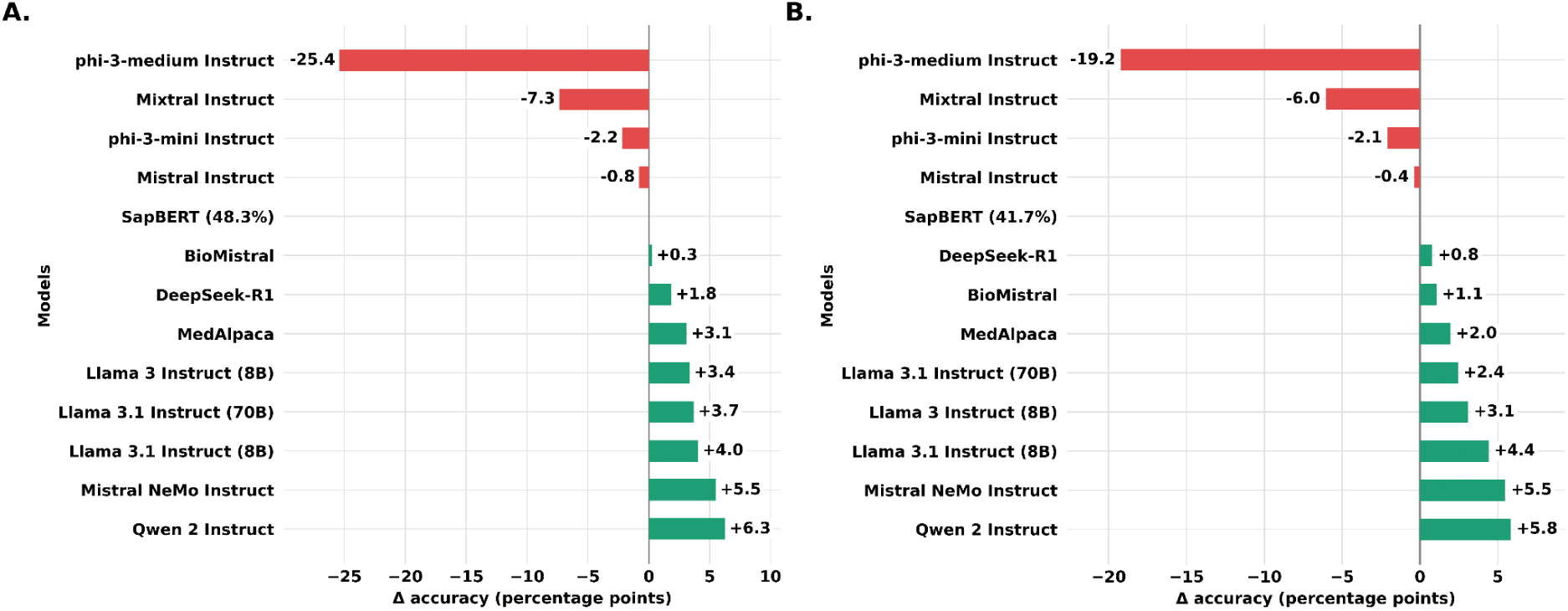
Effects of instruction-tuned LLM correction on Top-1 normalization accuracy relative to the standalone SapBERT baseline where positive values (green) represent improvement and negative values (red) represent performance degradation relative to the raw SapBERT pipeline **A)** SNOMED CT normalization and **B)** MedDRA normalization.

A similar pattern was observed for MedDRA normalization (Figure 6B). Relative to the standalone SapBERT baseline of 41.7%, Qwen 2 Instruct again produced the largest improvement (+5.8 pp), closely followed by Mistral NeMo Instruct (+5.5 pp). Llama 3.1 Instruct (8B), Llama 3 Instruct (8B), Llama 3.1 Instruct (70B), MedAlpaca, BioMistral and DeepSeek-R1 also improved on the baseline, whereas phi-3-medium Instruct (-19.2 pp), Mixtral Instruct (-6.0 pp), phi-3-mini Instruct (-2.1 pp) and Mistral Instruct (-0.4 pp) reduced Top-1 performance. Thus, the benefit of LLM-based correction was consistent across both vocabularies but remained strongly dependent on the choice of corrector.

The computational analysis in Supplementary Figures S10-S11 further highlights substantial differences in efficiency among the LLM correctors. For SNOMED CT, the mean end-to-end runtime over the full 12,713-query set across models was 3.94 hours (range: 2.01 - 8.11 hours), while for MedDRA it was 3.9 hours (range: 2.01 - 7.90 hours). Qwen 2 Instruct maintained a relatively high throughput of ∼1.53 - 1.55 queries per second, whereas Llama 3.1 Instruct (70B) processed only ∼0.76 - 0.78 queries per second, with Mistral Instruct, phi-3-medium Instruct and Mixtral Instruct showing still lower throughput. The largest model-dependent variation arose from the LLM correction stage, which averaged 2.11 hours for SNOMED CT and 2.09 hours for MedDRA, whereas FAISS index loading, retrieval and post-processing remained comparatively stable across models. These differences are consistent with the greater computational demands of larger or more complex models during autoregressive inference, making Qwen 2 Instruct a favorable compromise between normalization performance and computational efficiency. Furthermore, as illustrated in Table 4, integration of Qwen 2 Instruct with the SapBERT pipeline successfully corrected several false-negative predictions produced by standalone SapBERT, particularly for colloquial symptom descriptions, idiomatic/slang expressions and biomedical abbreviations, thereby improving alignment with the correct SNOMED CT and MedDRA concepts.

**Table 4:**
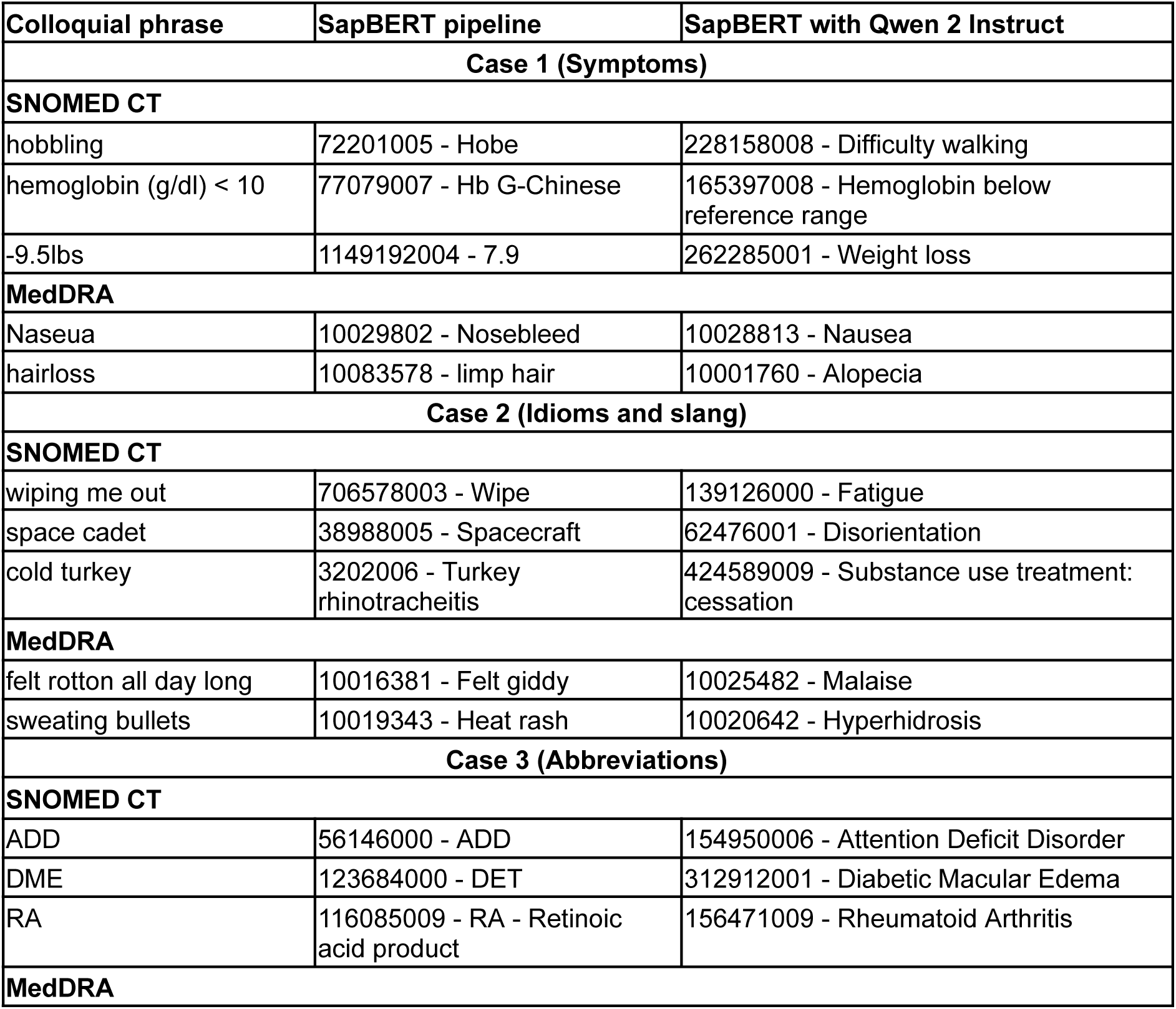
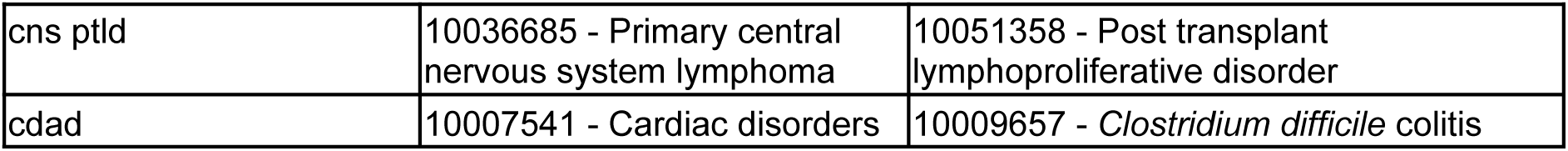
Evaluation-set instances that standalone SapBERT mapped incorrectly (false negatives) and that SapBERT with Qwen 2 Instruct mapped correctly (true positives), with their SNOMED CT and MedDRA concepts.

To further assess the robustness of the selected corrector, Qwen 2 Instruct was evaluated across ten repeated inference runs with randomized input ordering. Full-retrieval performance was completely invariant across runs, with a mean accuracy of 86.36%, 100% stable outcomes, 100% mean pairwise agreement and Cohen’s κ = 1. Top-1 performance showed minimal variability, with a mean accuracy of 54.63%, between-run SD of 0.033 percentage points and 99.53% stable outcomes, with no significant run effect (Cochran’s Q = 10.42, df = 9, P = 0.318). None of the 45 pairwise McNemar comparisons remained significant after Holm correction. Detailed reproducibility statistics are provided in Supplementary Tables S19 and S20.

## Semantic Organization and Nearest-Neighbor Retrieval

Figure 7 presents SapBERT (768-dimensional) embedding space visualization in two complementary panels. Figure 7A shows a 2D projection of UMLS term embedding space using all 127 semantic type unique identifiers (TUI), with each datapoint representing a term embedding colored consistently by TUI (n=8,998,517). This reveals large, dense and distinct clusters in which semantically related terms tend to form coherent regions rather than a wholly fragmented scatter, reflecting strong semantic alignment that underlies the superior normalization performance of the SapBERT model. Figure 7B illustrates a representative retrieval example, displaying the five nearest SNOMED CT term neighbors to the free-text query “*trouble with my liver*” within the SapBERT embedding space visualized in a 3D t-SNE projection. The top-ranked candidate was liver problems (TUI: T033; s=0.828), followed by Hepatic dysfunction NOS (TUI: T033; s=0.770), liver pain (TUI: T184; s=0.768), hepatic impairment (TUI: T047; s=0.754) and liver damage (TUI: T046; s=0.729). Despite spanning multiple semantic types (T033: Finding, T184: Sign or Symptom, T046: Pathologic Function and T047: Disease or Syndrome), all retrieved concepts remain clinically proximate to the query, demonstrating that SapBERT’s embedding space captures meaningful medical relationships that extend beyond strict semantic type boundaries, enabling top-ranked retrieval even from informal free-text expressions.

**Figure 7:**
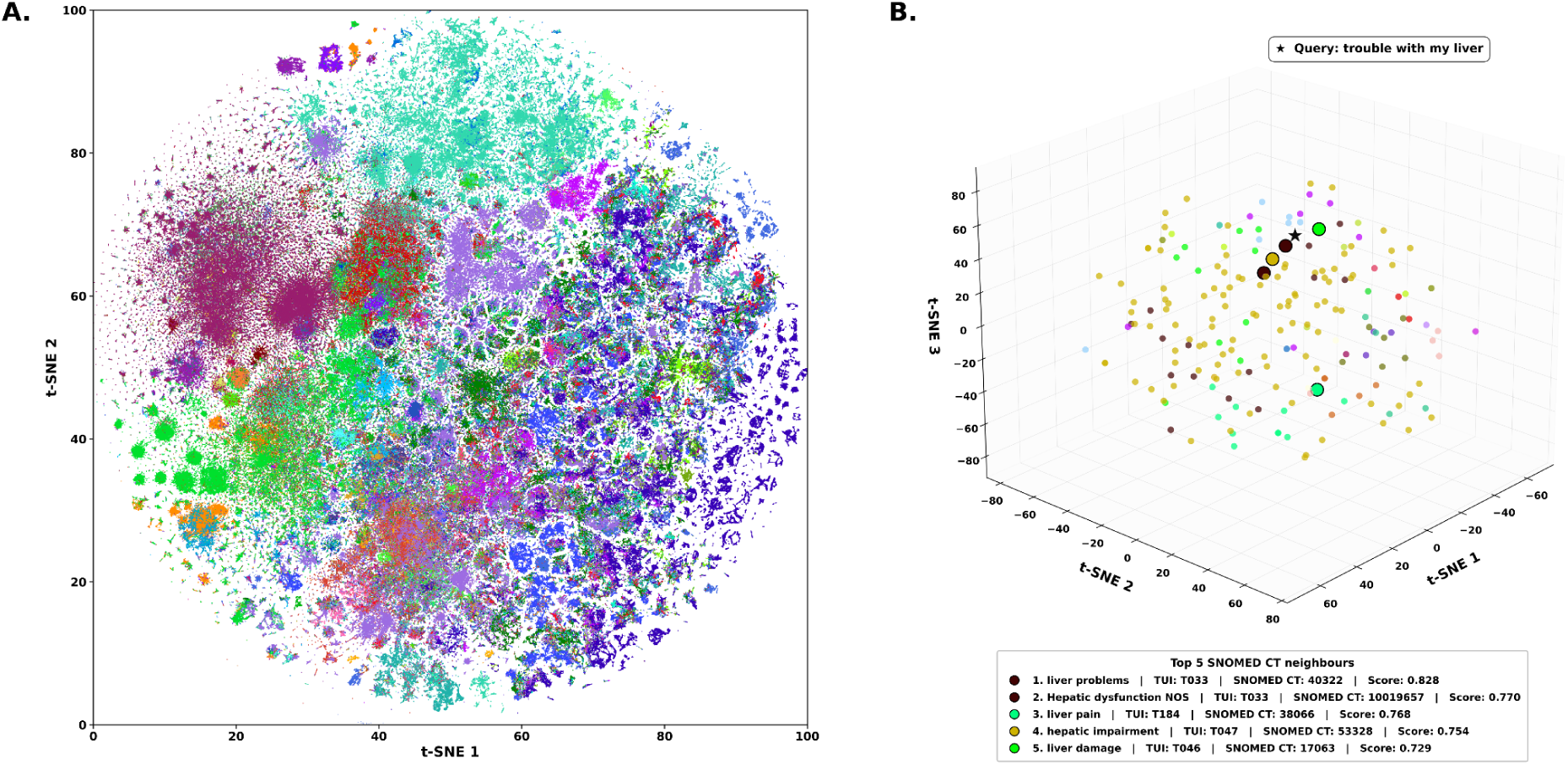
SapBERT embedding space visualization and nearest-neighbor retrieval. **A)** 2D t-SNE projection of the full SapBERT UMLS embedding space across all 127 active semantic types, with each datapoint representing a single UMLS term colored by semantic types (n=8,998,517). **B)** 3D t-SNE projection showing the five nearest UMLS term neighbors to the query “trouble with my liver”, with point sizes scaled by similarity score.

### Failure Analysis and Limitations

Supplementary Figure S2 shows that the models largely succeed on the same instances, implying a shared residue of difficult cases. Ultimately, even the Qwen 2 Instruct + SapBERT pipeline failed to normalize 1,734 instances for SNOMED CT and 2,933 instances for MedDRA, highlighting the persistent challenge of mapping noisy and ambiguous patient language.

Table 5 and Table 6 contain some of the instances for which our pipeline assigned wrong concepts. Table 5 highlights instances for which the ground truth does not align with the mapped medical codes from the Qwen 2 Instruct + SapBERT pipeline. Most of the errors were associated with semantic ambiguity, vague symptom descriptions or colloquial expressions that were mapped to related but clinically distinct concepts. For example, “*pain between my shoulder blades*” was mapped to “*Shoulder blade pain*”, whereas correct concepts were “*Back pain*” or “*Thoracic back pain*”. Similarly, a narrower mapping of *“Loose motions”* was produced instead of the broader concept of *“Diarrhoea”* for the phrase *“Loose stool”*.

**Table 5:** Representative retrieval-stage failure cases showing incorrectly mapped concepts against ground-truth concepts along with identifiers.

| Input phrase | Mapped entry | Ground truth |
| --- | --- | --- |
| **SNOMED CT** |  |  |
| pain between my shoulder blades | 20793008 - Shoulder blade pain | 161891005 - Back pain<br>279038004 - Thoracic back pain |
| head feels weird | 25064002 - Head pain | 271782001 - Drowsiness, Somnolence |
| gastric problems | 95516005 - Gastrointestinal upset | 29384001 - Stomach Diseases |
| hurts to get up in the morning | 40144003 - Morning stiffness | 22253000 - Pain<br>425423002 - Pain provoked by movement |
| shaky | 20262006 - Ataxia | 26079004 - Involuntary Quiver<br>267079009 - Trembling |
| **MedDRA** |  |  |
| Knee pain | 10064238 - Gonalgia | 10053156 - Musculoskeletal discomfort<br>10003239 - Arthralgia |
| Heart racing | 10043086 - Tachycardia, unspecified | 10033557 - Palpitations |
| Loose stool | 10024839 - Loose motions | 10012735 - Diarrhoea |
| eyes burn | 10006774 - Burning eyes | 10013774 - Dry eye |
| Ear infections | 10014011 - Ear infection | 10060945 - Bacterial infection |

**Table 6:** Representative correction-stage failure cases showing incorrect LLM-based pre-processing leading to incorrect mappings.

| Input phrase | LLM Correction | Ground truth |
| --- | --- | --- |
| **SNOMED CT** |  |  |
| afterpain | Postpartum pain | 22253000 - Pain |
| mtc | Muscle Tone Abnormality | 450886002 - Medullary Thyroid Carcinoma |
| Coxalgia | Lower back pain | 57676002 - Joint pain |
| lucidity | Clarity | 85418005 - Dream disorder |
| malaisse | Fatigue | 367391008 - Malaise |
| **MedDRA** |  |  |
| shroke | Stridor | 10008190 - Cerebrovascular accident |
| Logorrhoea | Excessive watery secretion from the mouth | 10024796 - Logorrhoea |
| Otosalpingitis | Otitis media with mastoid involvement | 10033102 - Otosalpingitis |
| rpls | Replaced | 10071066 - Posterior Reversible Encephalopathy Syndrome |
| direar | Ear pain | 10012735 - Diarrhoea |

Although the overall results show considerable improvement over the baseline, there are cases where the LLMs become a source of noise themselves, introducing subtle semantic drifts. A few such instances are reported in Table 6, where, for example, the *“mtc”* acronym was incorrectly expanded to *“Muscle Tone Abnormality”,* driving the normalization astray from the standardized terminology *“Medullary Thyroid Carcinoma”*. Similarly, anatomical shifts such as the supposed correction of *“Coxalgia”* to *“Lower back pain”* again result in a failure case. Over-generalization has the same effect: the LLM rendered "malaisse" as "Fatigue" instead of correcting the spelling to "malaise", leaving the query unmapped to its gold-standard code. These observations indicate that failures arise not only from retrieval mismatches but also from contextually incorrect LLM reformulations upstream. Such confidently wrong, structurally plausible failure modes require LLM-mediated clinical tools to be deployed cautiously.

As summarized in Table 7, false-negative instances predominantly reflected lexical-only, clinically safe reformulations, whereas a smaller subset involved clinically meaningful errors, including concept substitution, incorrect acronym expansion, altered specificity and anatomical shifts. Additional errors arose from moderately severe transformations and formatting or Unicode incompatibilities.

**Table 7:** Clinical characterization of LLM reformulations among false-negative normalization instances.

| No. | Reformulation category | SNOMED CT (n) | MedDRA (n) |
| --- | --- | --- | --- |
| **1** | Clinically critical incorrect reformulation | 402 | 461 |
| 1(a) | Concept substitution | 327 | 373 |
| 1(b) | Incorrect acronym expansion | 34 | 34 |
| 1(c) | Over-generalization/Over-specialization | 37 | 49 |
| 1(d) | Anatomical shift | 4 | 5 |
| **2** | Moderately severe transformation | 156 | 204 |
| **3** | Formatting and Unicode incompatibilities | 125 | 156 |
| **4** | Total false-negative instances | 1,734 | 2,933 |
|  | Lexical-only, clinically safe transformations | 1,051 | 2,112 |

This study has several limitations. First, evaluation was limited to the SNOMED CT and MedDRA as ground-truth vocabularies from MedNorm, as no suitable annotated datasets were found for other vocabularies. Second, although bootstrap confidence intervals were incorporated for the Qwen reproducibility analysis, they were not systematically computed across all transformer-based embedding models and instruction-tuned LLMs because of computational and time constraints. Third, the framework was evaluated on isolated words and short phrases and not full-fledged sentences or EHR documents. Direct comparison with established MCN systems was also not performed because of substantial differences in task design, supervision and evaluation settings; therefore, performance claims are limited to the models benchmarked under the same zero-shot retrieval framework. Finally, this framework lacks essential safeguards such as confidence-based abstention, uncertainty calibration, human review and drift detection, which are required for critical clinical use in the real world.

## Conclusion

In this study, we developed a semantic mapping-based pipeline for MCN of loose, informal and user-generated biomedical text. The framework integrates an Instruction-tuned LLM corrector with SapBERT encoding and a UMLS-derived semantic space indexed via FAISS for efficient cosine similarity retrieval. SapBERT delivered the strongest standalone results, while adding an LLM correction layer further improved handling of linguistically irregular inputs. Llama 3.1 Instruct (70B) achieved the highest overall accuracy, whereas Qwen 2 Instruct provided a better trade-off between performance and computational efficiency. Notably, the largest gains occurred on the highly noisy SMM4H2017 dataset; improvements on cleaner benchmarks were more variable, demonstrating that correction is most useful for fragmented, colloquial and misspelled expressions. These findings highlight the value of combining LLM-based text normalization with ontology-aware embedding retrieval, offering a scalable and practical solution for real-world biomedical concept mapping tasks.

## Key Points

1. The proposed framework requires no manual annotation, operates without named entity recognition (NER) and accepts grammar-agnostic input, thereby eliminating pre-processing overhead and enabling processing of raw biomedical text.

2. The system is grounded in the Unified Medical Language System (UMLS), providing vocabulary-agnostic coverage that extends beyond any single lexicon dataset and spans the full UMLS concept universe.

3. To the best of our knowledge, this is the first study to construct and systematically compare the complete UMLS concept embedding spaces using 15 general, biomedical and clinical language models, enabling large-scale evaluation of their ability to capture and organize biomedical semantic relationships.

4. Concept normalization is performed at the term-level using metric learning models, enabling rich synonym representation and transcending surface form limitations inherent to lexicon-based approaches.

5. The end-to-end pipeline is fully automated, producing standardized structured output from raw input without any human intervention, making it readily scalable for large-scale biomedical text mining applications.

## Supporting information

Source Data for Supplementary Figures S4-S9

## Acknowledgements

The authors thank the IT Division of CSIR-IGIB and CSIR-NIDSA (formerly CSIR-4PI) for infrastructure and computational facilities. AV acknowledges IndiaAI for a fellowship. A, MV and SP acknowledge CSIR for their fellowships.

## Author Contributions

KC, AV and SP conceptualized the study. AV and A curated the data and performed the formal analysis. AV and A were responsible for investigations presented within the study. AV, A and KC designed the methodology. KC was responsible for project administration. AV and A were responsible for software. KC, AV and SP were responsible for overall supervision. AV, A, MV and SP performed data validation. AV, A and MV were responsible for visualization. AV and A wrote the original draft. KC, AV, A, MV and SP reviewed and edited the manuscript. All authors approved the final draft of the manuscript.

## Supplemental Material

Supplementary material is provided separately. Supplementary Figures S1-S11 and Supplementary Tables S1-S2 are provided in Supplementary Information. Supplementary Tables S3-S20, containing all statistical results underlying the reported comparisons, are provided as separate excel workbooks. Source data for Figures 3-5 and Supplementary Figures S4-S9 are provided in Source Data for Supplementary Figures S4-S9.

## Data and Code Availability

All data used in this study is publicly available. The code supporting the findings of this study is currently hosted in a private repository and will be made publicly available upon acceptance at https://github.com/DiGeMed/MedLexAlign.

