## Supplementary Information for "A Comparative Benchmark of Biomedical Language Models for Concept Normalization from Real-World Text"

***Equal First Author**

**^#^Corresponding Author**

**Affiliations:**

^1^CSIR-Institute of Genomics and Integrative Biology, New Delhi, 110007 India

^2^Academy of Scientific and Innovative Research (AcSIR), Ghaziabad, 201002 India

^3^Amity Institute of Biotechnology, Amity University, Noida, 201313 India

**Supplementary Material**

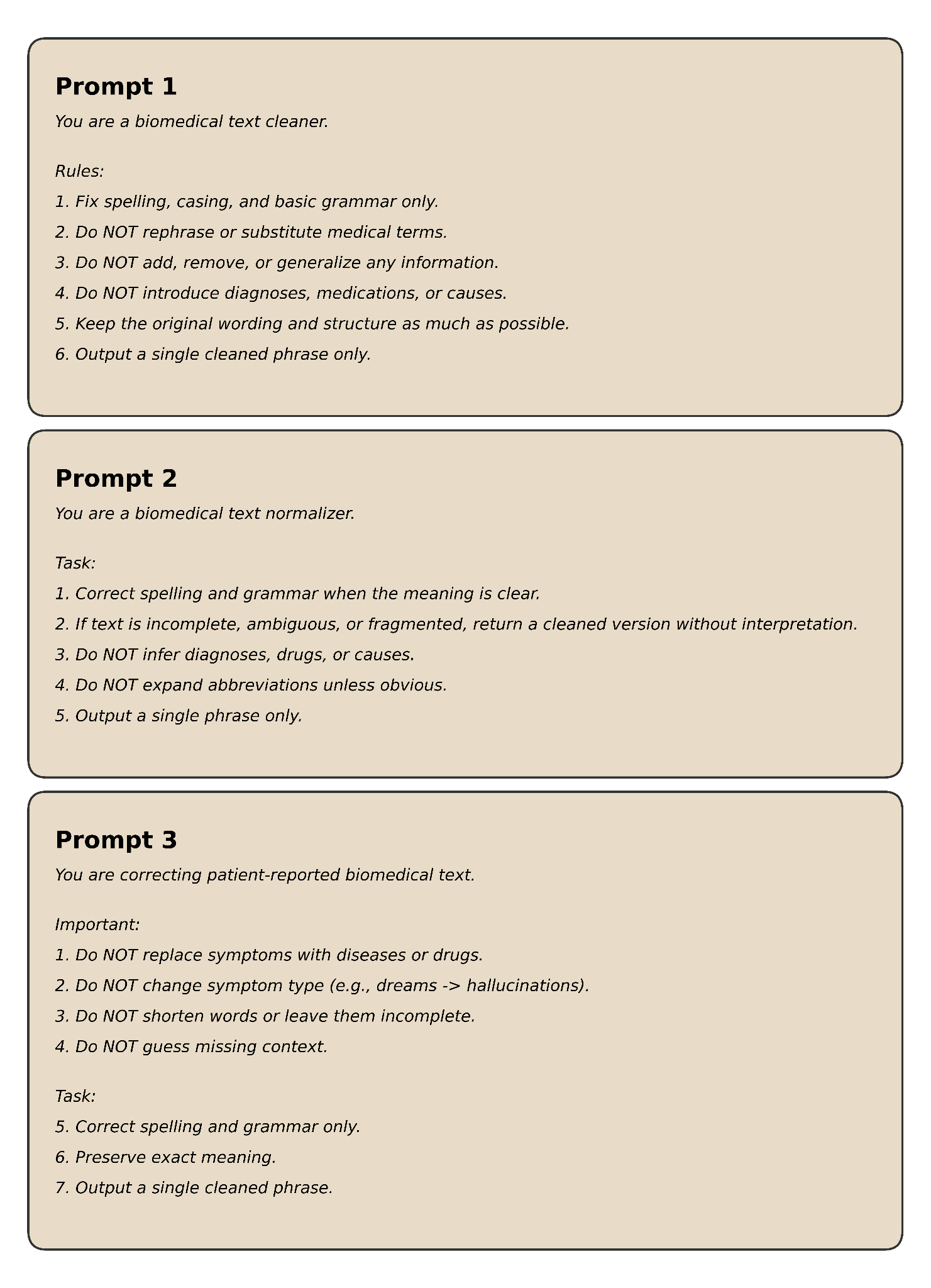

**Supplementary Figure S1:** The three remaining specialized prompt templates (Prompts 1 – 3) evaluated for query correction; the selected template (Prompt 4) is shown in Figure 2.

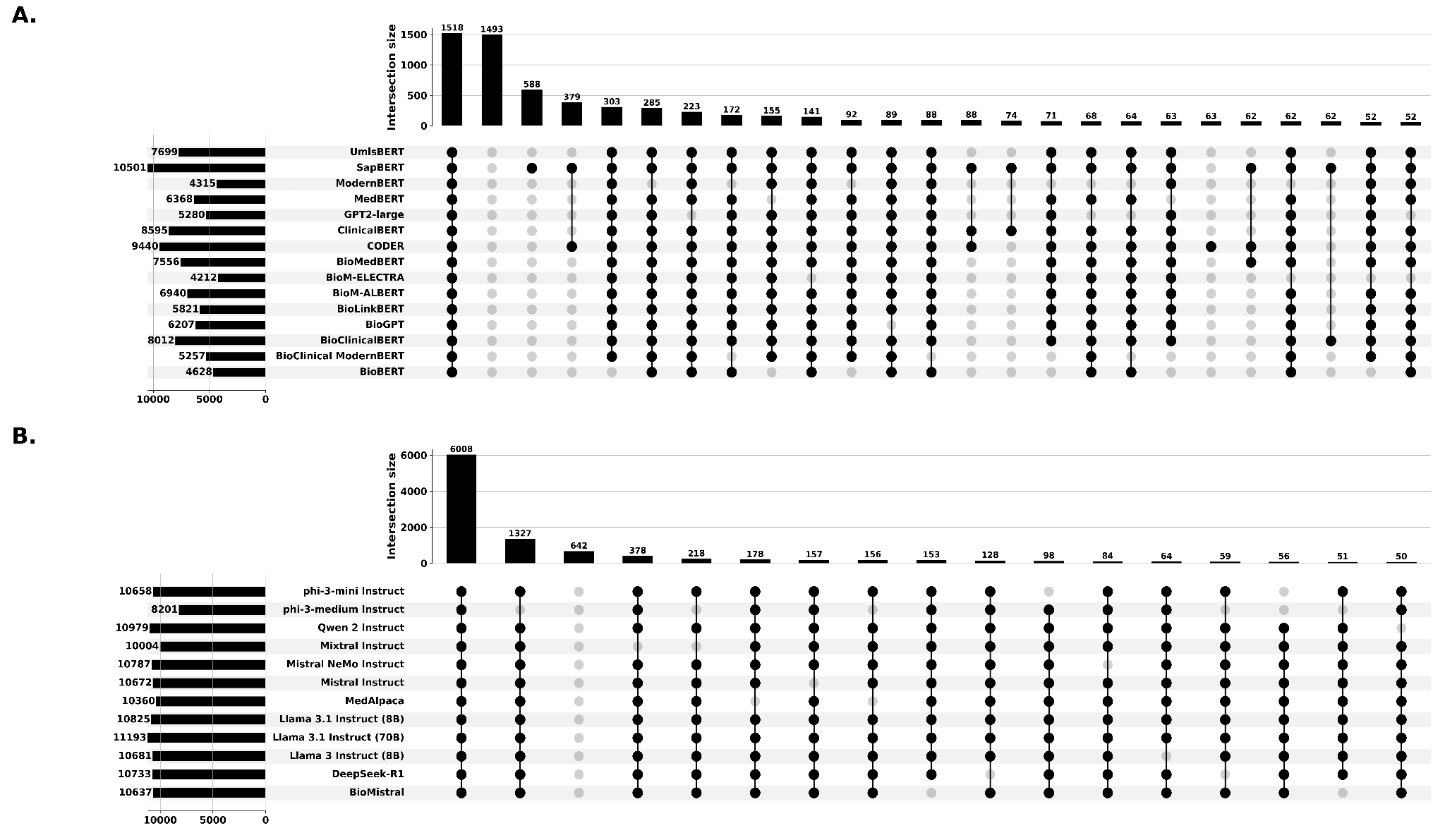

**Supplementary Figure S2:** UpSet plot illustrating overlaps in correctly normalized instances across models on the blind benchmark datasets for **A)** transformer-based models and **B)** LLM-based pipelines. Left bars show the total correctly mapped instances per model, while top bars show exact intersection sizes. Connected black dots indicate models included in each intersection; only intersections with at least 50 instances are shown.

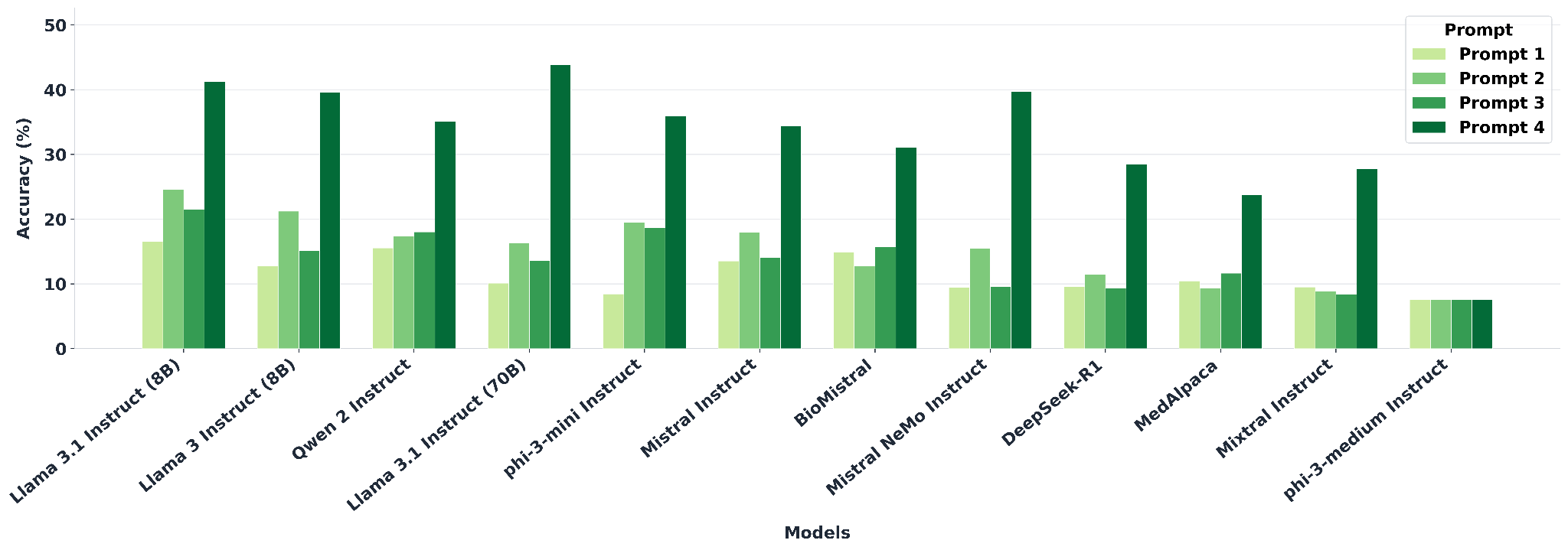

**Supplementary Figure S3:** Comparative performance of four candidate prompt designs evaluated on the challenging instances that were not correctly normalized by any of the transformer-based models. The best-performing prompt (Prompt 4), based on improved correction and downstream mapping accuracy was selected for all subsequent LLM-assisted correction experiments.

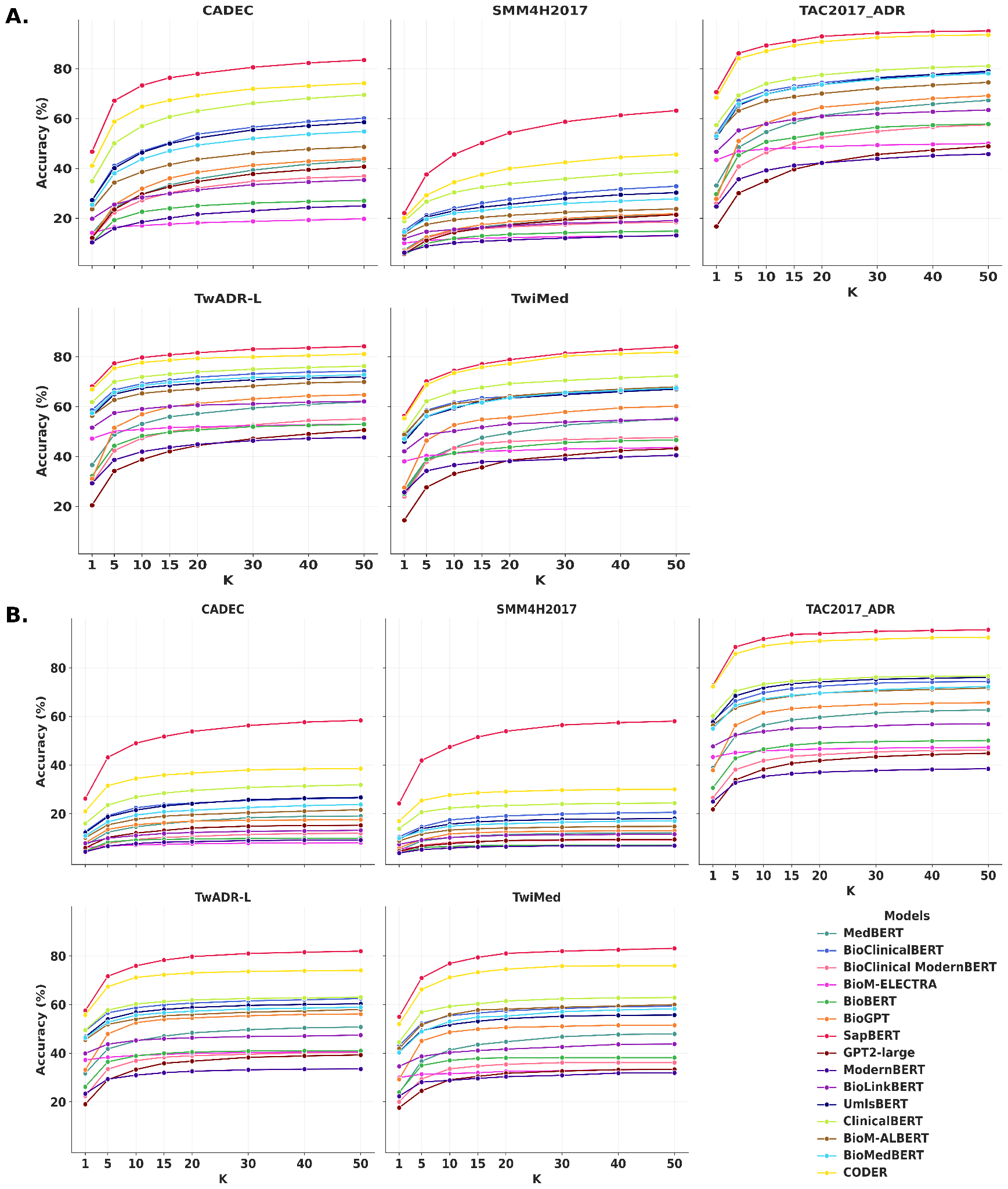

**Supplementary Figure S4:** Top-K retrieval accuracy curves for each dataset, depicting the proportion of correct concept matches within the Top-K candidates (K = 1 – 50) across transformer-based models. **A)** SNOMED CT-based evaluation and **B)** MedDRA-based evaluation.

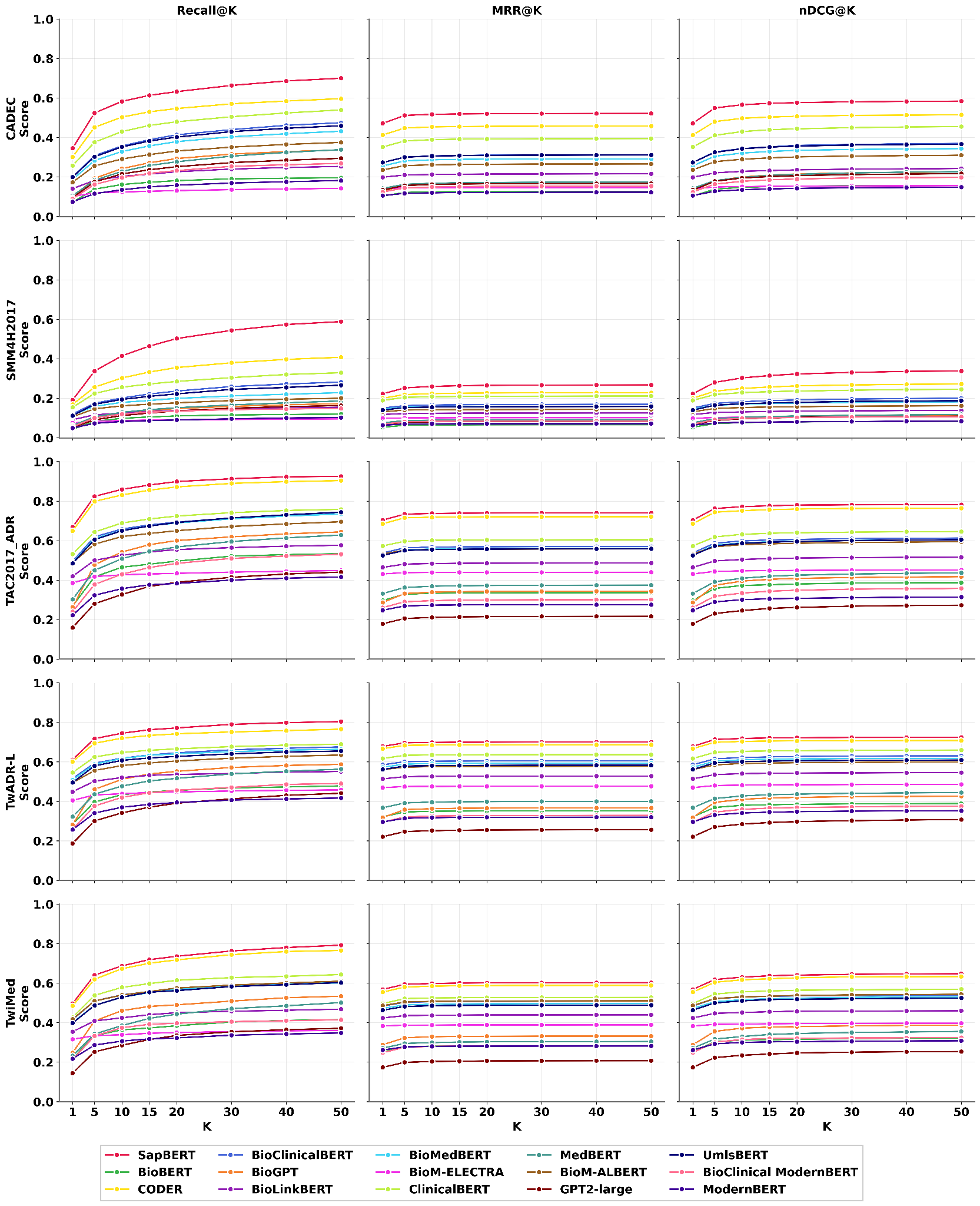

**Supplementary Figure S5:** Recall@K, Mean Reciprocal Rank (MRR@K) and Normalized Discounted Cumulative Gain (nDCG@K) are shown for all transformer-based models across the five benchmark datasets for SNOMED CT.

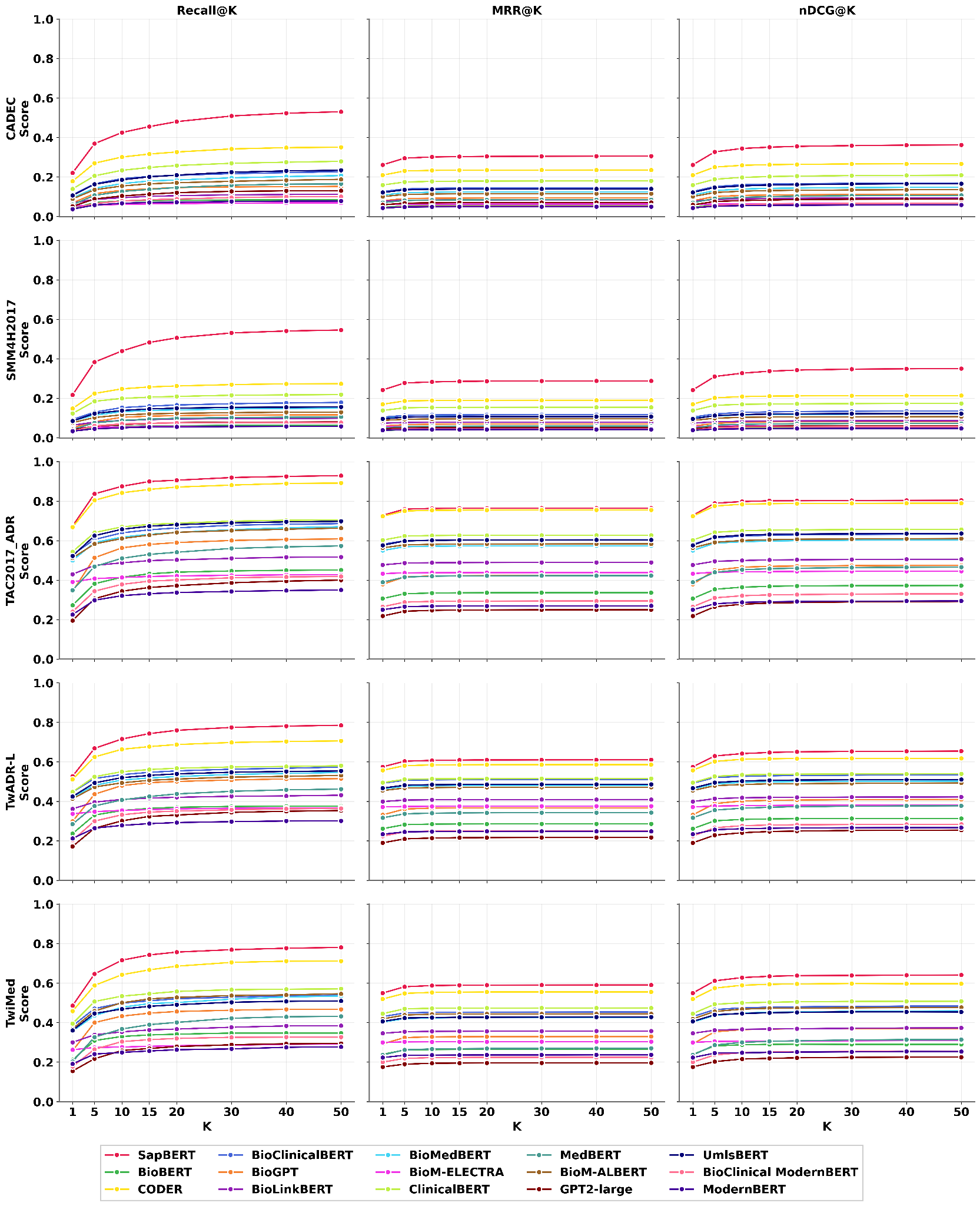

**Supplementary Figure S6:** Recall@K, Mean Reciprocal Rank (MRR@K) and Normalized Discounted Cumulative Gain (nDCG@K) are shown for all transformer-based models across the five benchmark datasets for MedDRA.

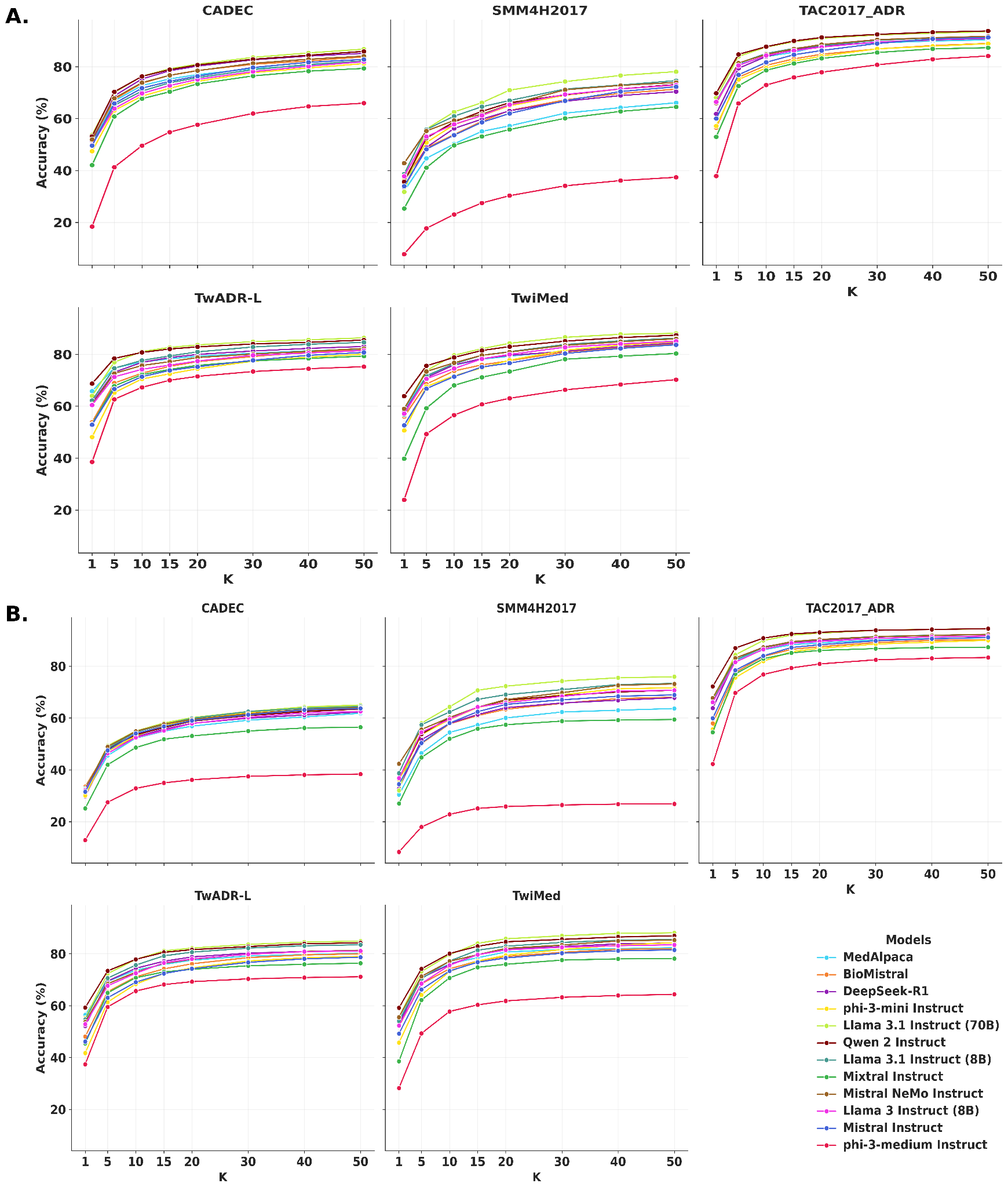

**Supplementary Figure S7:** Top-K retrieval accuracy curves for each dataset, depicting the proportion of correct concept matches within the Top-K candidates (K = 1 – 50) across instruction-tuned models along with SapBERT. **A)** SNOMED CT-based evaluation and **B)** MedDRA-based evaluation.

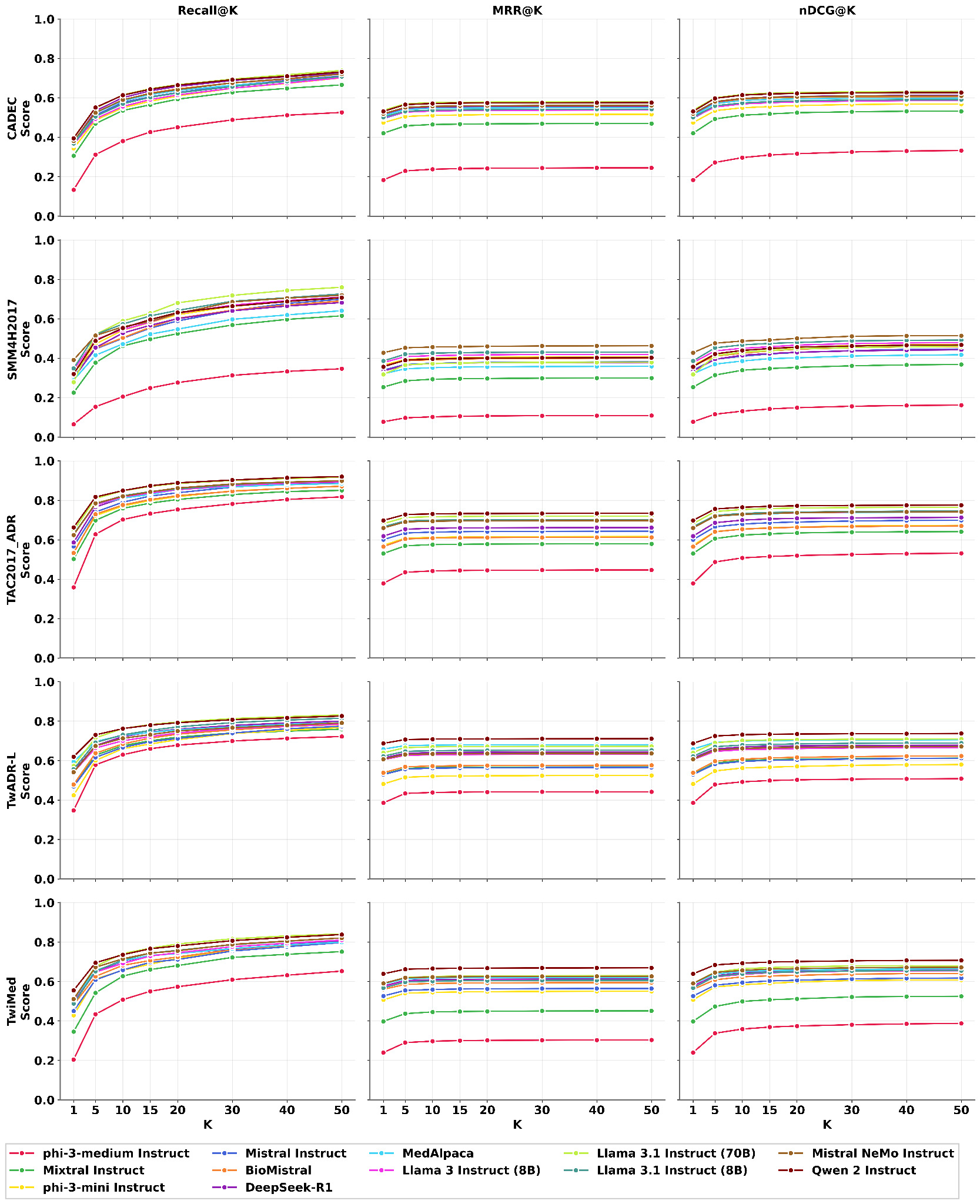

**Supplementary Figure S8:** Recall@K, Mean Reciprocal Rank (MRR@K) and Normalized Discounted Cumulative Gain (nDCG@K) are shown for all instruction-tuned models along with SapBERT across the five benchmark datasets for SNOMED CT.

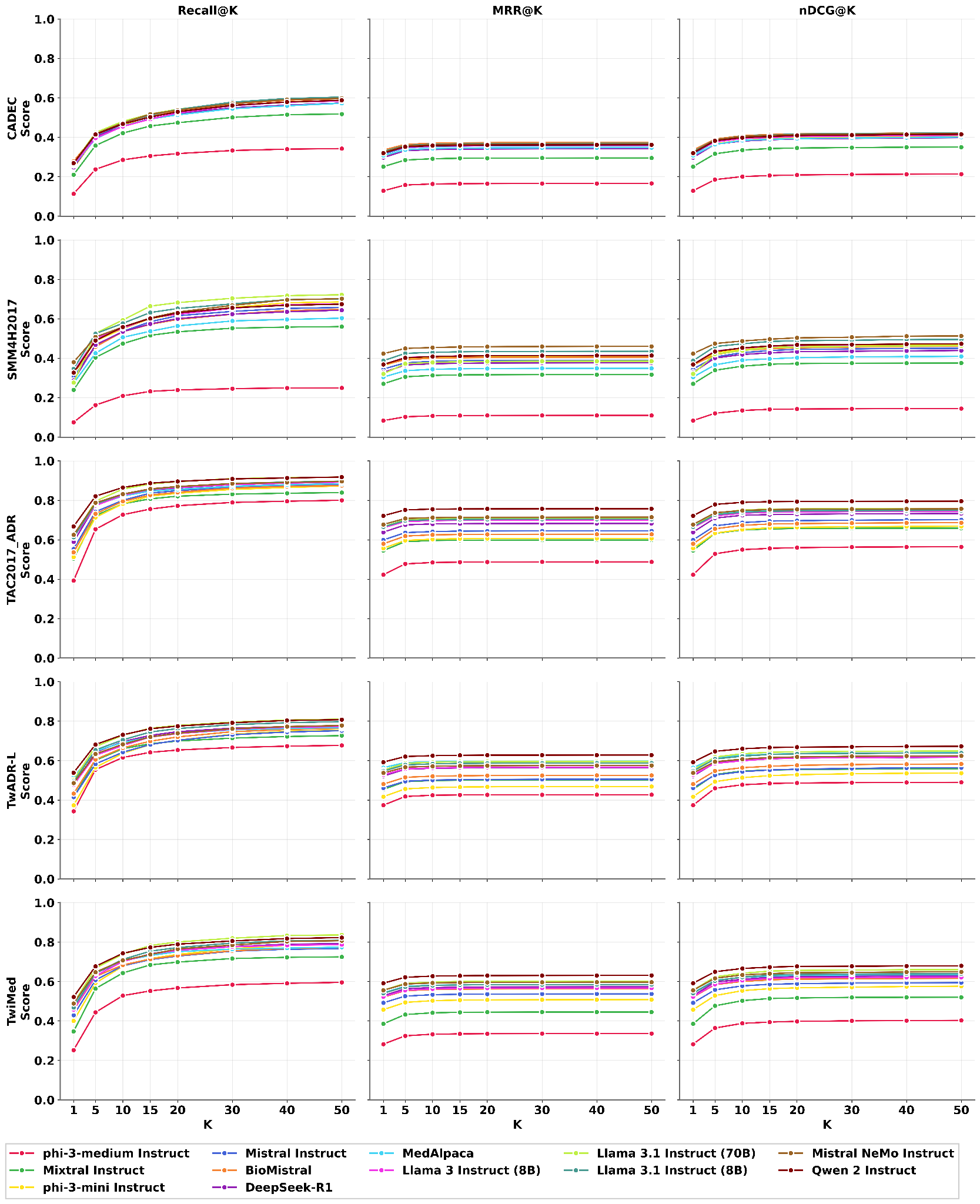

**Supplementary Figure S9:** Recall@K, Mean Reciprocal Rank (MRR@K) and Normalized Discounted Cumulative Gain (nDCG@K) are shown for all instruction-tuned models along with SapBERT across the five benchmark datasets for MedDRA.

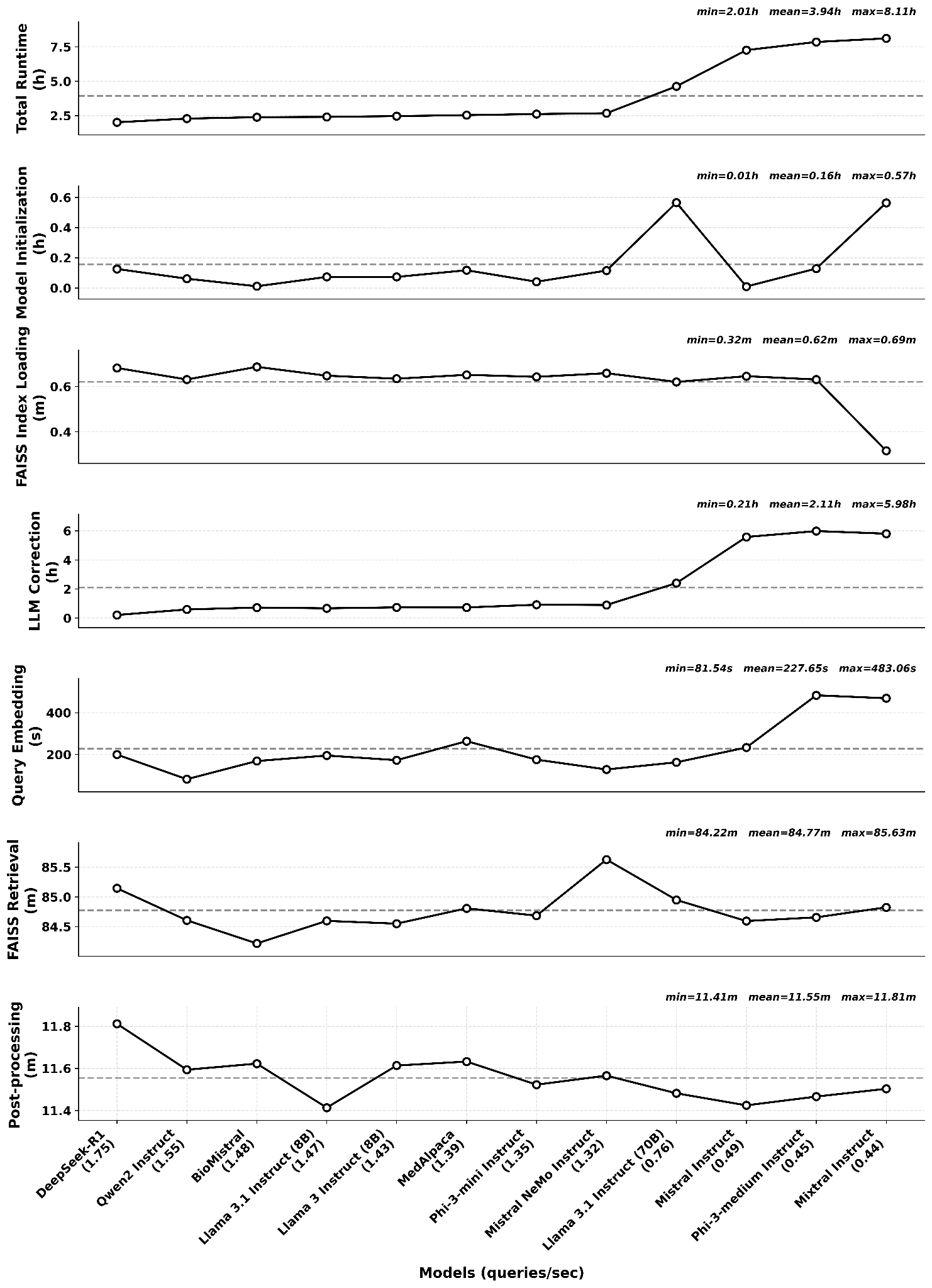

**Supplementary Figure S10:** Runtime comparison of LLM-assisted medical concept normalization models across different stages: total runtime, model initialization, LLM correction, query embedding, FAISS retrieval and post-processing for SNOMED CT over the full 12,713-query evaluation set.

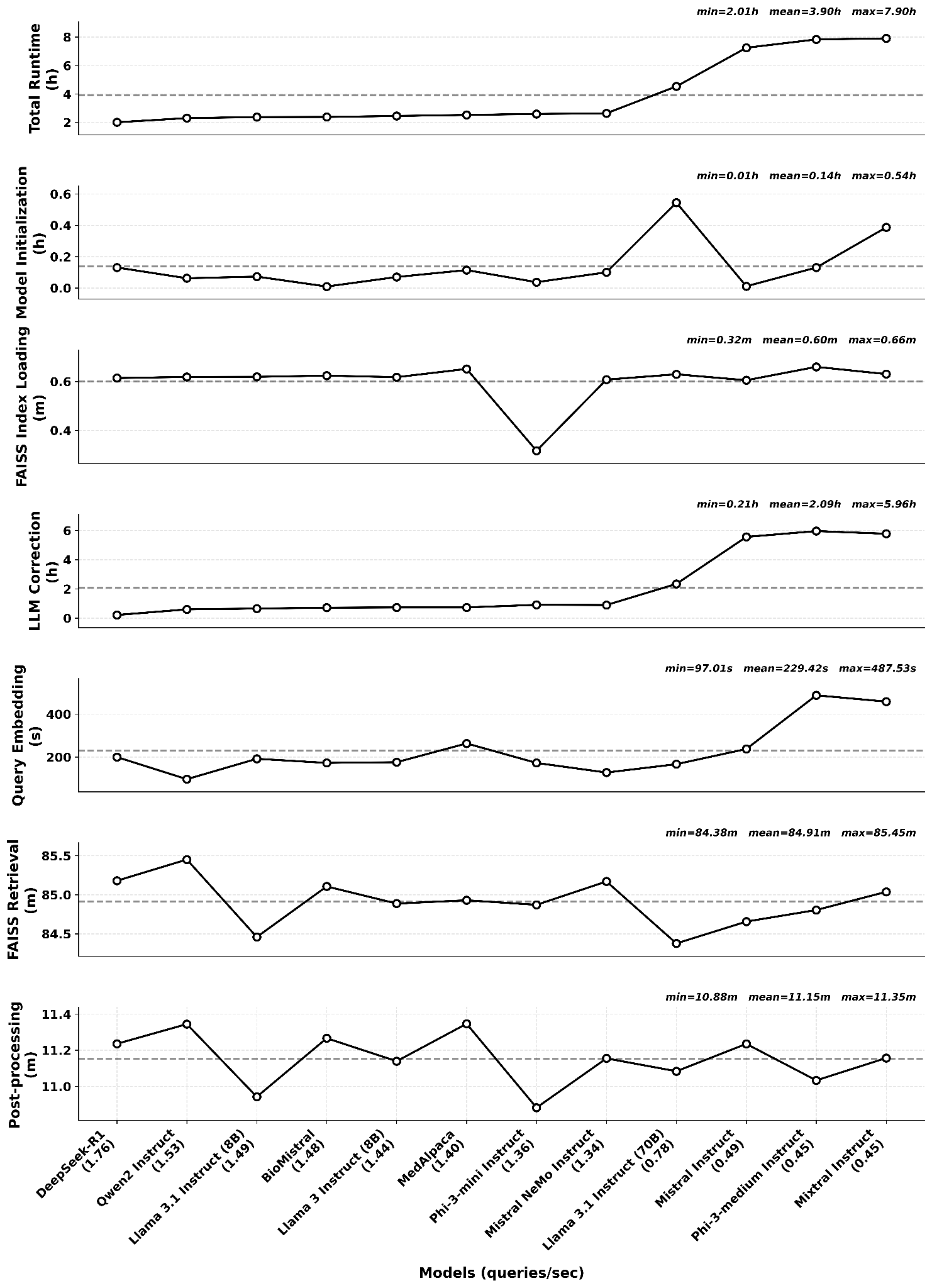

**Supplementary Figure S11:** Runtime comparison of LLM-assisted medical concept normalization models across different stages: total runtime, model initialization, LLM correction, query embedding, FAISS retrieval and post-processing for MedDRA over the full 12,713-query evaluation set.

**Supplementary Table S1:** Top source abbreviation (SABs) included in the indexed UMLS subset, showing vocabulary abbreviation, term count, percentage contribution and expanded terminology name.

| **Vocabulary** | **Count** | **Percentage (%)** | **Expanded name** |
| --- | --- | --- | --- |
| **SNOMEDCT_US** | 1,688,367 | 16.10 | Systemized Nomenclature of Medicine-Clinical Terms |
| **MEDCIN** | 1,055,425 | 10.06 | MEDCIN clinical terminology |
| **NCBI** | 1,043,954 | 9.95 | National Center for Biotechnology Information |
| **MSH** | 1,030,981 | 9.83 | Medical Subject Headings |
| **LNC** | 742,614 | 7.08 | Logical Observation Identifiers Names and Codes |
| **NCI** | 488,231 | 4.65 | National Cancer Institute Thesaurus |
| **RXNORM** | 354,029 | 3.37 | Prescription Normalization |
| **ICD10PCS** | 351,821 | 3.35 | International Classification of Diseases, 10th Revision - Procedure Coding System |
| **RCD** | 347,568 | 3.31 | Read Codes |
| **MTH** | 259,444 | 2.47 | Metathesaurus - internal UMLS Metathesaurus source entries |
| **HGNC** | 236,374 | 2.25 | HUGO Gene Nomenclature Committee |
| **MTHSPL** | 211,702 | 2.02 | Metathesaurus Supplemental - UMLS supplemental Metathesaurus entries |
| **ICD10CM** | 208,979 | 1.99 | International Classification of Diseases, 10th Revision - Clinical Modification |
| **OMIM** | 200,948 | 1.92 | Online Mendelian Inheritance in Man |
| **GO** | 181,225 | 1.73 | Gene Ontology |

**Supplementary Table S2:** Overview of the MedNorm evaluation dataset used for blind evaluation, summarizing phrase counts and corresponding SNOMED CT ,MedDRA and UMLS identifiers across constituent datasets before removing redundancy.

| **Dataset** | **Phrases** | **SNOMED IDs** | **MedDRA IDs** | **UMLS CUIs** |
| --- | --- | --- | --- | --- |
| **TAC2017_ADR** | 5,835 | 5,835 | 5,835 | 0 |
| **TwADR-L** | 4,626 | 4,626 | 4,626 | 4,626 |
| **TwiMed** | 1,858 | 1,858 | 1,858 | 1,858 |
| **CADEC** | 6,797 | 6,797 | 6,797 | 0 |
| **SMM4H2017** | 8,863 | 8,863 | 8,863 | 0 |
| **Total** | 27,979 | | | 6,484 |
